# Resolving forebrain developmental organisation by analysis of differential growth patterns

**DOI:** 10.1101/2025.01.10.632351

**Authors:** Elizabeth Manning, Kavitha Chinnaiya, Caitlyn Furley, Dong Won Kim, Seth Blackshaw, Marysia Placzek, Elsie Place

## Abstract

The forebrain is the most complex region of the vertebrate CNS, and its developmental organisation is controversial. We fate-mapped the embryonic chick forebrain using lipophilic dyes and Cre-recombination lineage tracing, and built a 4D model of brain growth. We reveal modular patterns of anisotropic growth, ascribed to progenitor regions through multiplex HCR. Morphogenesis is dominated by directional growth towards the eye, more isometric expansion of the prethalamus and dorsal telencephalon, and anterior movement of ventral cells into the hypothalamus. Fate conversion experiments in chick and comparative gene expression analysis in chick and mouse support placement of the hypothalamus ventral to structures extending from the telencephalon up to and including the zona limitans intrathalamica (ZLI), with the dorsoventral axis becoming distorted at the base of the ZLI. Our findings challenge the widely accepted prosomere model of forebrain organisation, and we propose an alternative ‘tripartite hypothalamus’ model.

---

In recent years, remarkable progress has been made in the profiling and classification of central nervous system cell types, and in understanding their specification and differentiation from neural progenitors (*1–3*). Nonetheless, the fundamental layout of the developing vertebrate brain is still disputed, reflected in a confused and contradictory contemporary literature, with the extent and position of the hypothalamus at the centre of the debate. In classic columnar models of forebrain organisation, the thalamus, prethalamus, hypothalamus and eye were held to belong to the diencephalon, with the telencephalon positioned anterior to these (*4*, *5*). However, an alternative view, in the form of the ‘prosomere’ model, has since gained significant traction. In this scheme, the anteroposterior (A-P) and dorsoventral (D-V) axes are reinterpreted, and the forebrain consists of five transverse segments – ‘prosomeres’ – analogous to the neuromeres of the hindbrain and spinal cord. The prosomere model suggests that the hypothalamus and eye share the segmental identity of the telencephalon, lying anterior to the prethalamus and other diencephalic structures (*4*, *6*). However, the prosomere model has been challenged by recent studies showing that hypothalamic patterning is closely coordinated with prethalamic, rather than telencephalic, patterning (*7–11*).

Here we clarify forebrain developmental organisation. We completed the fate map for the Hamburger-Hamilton stage 10 (HH10) chicken anterior neural tube while systematically visualising growth patterns, enabling a 4D reconstruction of forebrain growth. DiI/DiO ‘growth lines’ reveal how region-specific patterns of anisotropic growth sculpt the amniote forebrain. Fate conversion experiments and molecular studies of the early chicken and mouse forebrain contradict key claims of the prosomere model. Instead our data indicates a central role of eye morphogenesis in separating the telencephalon from the prethalamus and hypothalamus, all within a common anterior forebrain compartment. Further, our data reveals a contiguous transverse boundary region, encompassing the supramammillary hypothalamus and *zona limitans intrathalamica* (ZLI), that divides the forebrain; region-specific growth distorts this transverse morphology, creating a flexure in the D-V axis at the base of the ZLI. Our work resolves prior misinterpretation and provides a new model for forebrain developmental organisation that we term the ‘tripartite hypothalamus’ model.

## Results

### DiI/DiO ‘growth lines’ reveal forebrain anisotropic growth

We fate-mapped the chicken forebrain at HH10, aided by prominent morphological landmarks including a series of transient epithelial folds in the ventral midline (Fig. 1A-C, fig. S1A-D). Embryos were injected with DiI/DiO, targeting the twelve areas depicted in Fig. 1D, and analysed at HH18-20. We labelled relatively large areas to provide insight into tissue-level growth patterns, and describe the resulting DiI/DiO distributions on the basis of morphology and marker genes (Fig. 1E, fig. S1E) (*7*, *12*). For descriptive purposes, we define the A-P axis as running parallel to the hypothalamic *NKX2-2* expression domain and the D-V axis as orthogonal to it (Fig. 1F).

**Figure 1:**
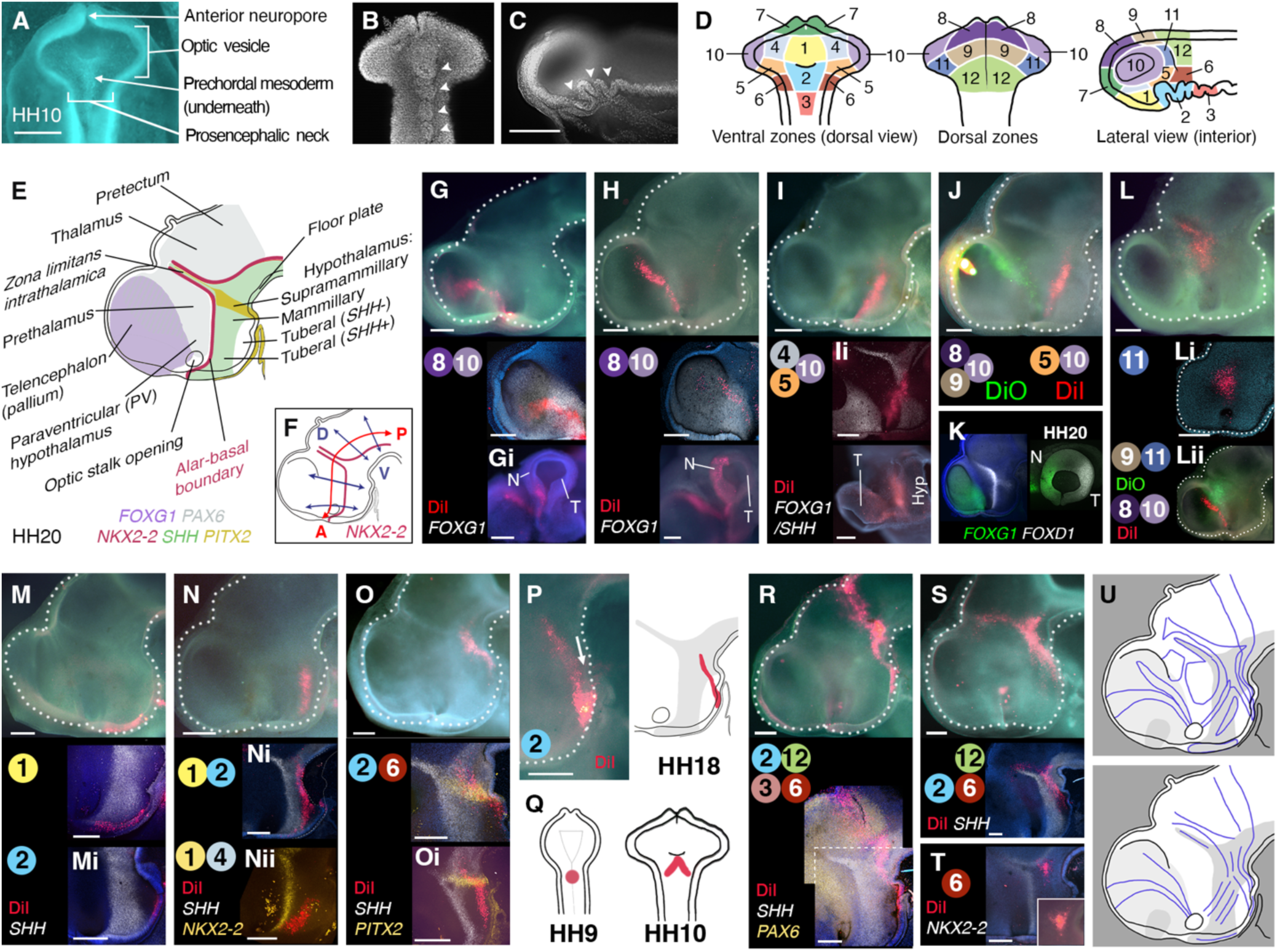
Fate mapping reveals forebrain growth patterns. **A-C)** HH10 chick morphology. **A)** *In ovo* dorsal view, dorsal neuroectoderm removed. **B)** DAPI-stained neuroepithelium, ventral view. **C)** hemisected, internal view. Arrowheads = midline folds. **D)** HH10 injection areas. **E)** HH20 forebrain regions and marker genes. **F)** A-P/D-V axes. **G-J,L-P,R-S)** Hemisected HH18-20 heads showing ‘growth lines’, DiI/DiO injected at HH10, internal views. Circled numbers indicate HH10 injection areas in (D); multiple numbers indicate intersection points (G-O) or multiple injections (R-S). Upper image: brightfield plus DiI/DiO. Lower: HCR *in situ* results for selected genes as indicated, and alternative views; same sample unless indicated by i, ii. Eye images are flipped horizontally to aid interpretation. **K)** HCR-processed internal and outer eye view of HH20. **Q)** Posterior ventral midline injection at HH9 forms V-shapes at HH10 and HH18. **U)** Summary of growth lines (upper) emphasising directionality (lower). n = 14-49 examples per pattern, total number of analysed growth lines (HH10 injections) = 225. Hyp - hypothalamus, N - Nasal eye, T - Temporal eye. Scale bars - 250 μm.

Labelled neuroectoderm reproducibly gave rise to distinctively shaped territories, referred to as ‘growth lines’, revealing highly anisotropic region-specific growth patterns (fig. S2). Growth lines originating in area 8 stretch from the *FOXG1*^+ve^ dorsal telencephalon through the dorsal optic stalk, terminating in the peripheral nasal retina (Fig. 1G-H). Growth lines arising from area 4/5 stretch from the *SHH*^+ve^ posterior hypothalamus, across the alar-basal boundary (ABB), into the ventral optic stalk and temporal retina (Fig. 1I-J). These long growth lines tend to stay within either *FOXG1^+ve^* or *FOXD1^+ve^*domains, and outline a wedge-shaped area of more isometric growth encompassing the prethalamus and paraventricular (PV) hypothalamus, which originate in area 11 (Fig. 1K-L).

Cells originating in areas 1-3 maintain their relative A-P positions near the ventral midline, contributing to the anterior *SHH^+ve^*(area 1, Fig. 1M) and *SHH^-ve^* tuberal hypothalamus, mammillary (MM) hypothalamus, and diencephalic floor plate (FP) (fig. S3A-B) (*12–14*). More lateral parts of areas 1, 2, 4, 5 (not previously fate-mapped) contribute to the *SHH^+ve^* and *SHH^-ve^* basal hypothalamus (Fig 1N). Long growth lines in the tuberal hypothalamus orientate in an A-P direction and extend towards the eye (Fig. 1N). Growth lines arising from areas 2/6 slope anteroventrally towards the tuberal hypothalamic midline, curving around the *FOXA1/2^+ve^* FP and becoming distorted as they cross the *PITX2^+ve^* supramammillary (SM) hypothalamus (Fig. 1O). Wider midline injections form an anteroventrally-sloping V-shape, showing that midline cells are displaced anteriorly relative to their more lateral counterparts (Fig. 1P, white arrow; fig. S3C-E, Fig. 1Q).

In contrast to the A-P growth lines in the basal hypothalamus, those in the thalamus and pretectum align in a D-V direction (Fig.S3F, area 12). The orientation of growth lines changes abruptly close to the ABB, just posterior to the ZLI (Fig. 1R-S). A small tricorn formation arose at this point (Fig. 1T, area 6). In summary, morphogenesis of the ventral anterior forebrain occurs in a stereotyped pattern of anisotropic growth, manifest as a series of overlapping growth lines (summarised in Fig. 1U). Key growth patterns, including the V-shaped and A-P oriented lines in the basal hypothalamus, and D-V oriented lines in dorsal regions posterior to the ZLI, were confirmed by DiI-DiO labelling of HH7 neuroectoderm (fig. S4), by genetic clonal analysis using the Cre-Cytbow transgenic line (*15*), and in RFP-electroporated embryos (fig. S5).

### Region-specific growth patterns describe forebrain morphogenesis

We built a 4-dimensional model of forebrain growth, based on the above findings combined with existing fate maps for dorsal regions (*16–18*) (fig. S6). This provides the most complete fate map yet of the HH10 chicken forebrain (Fig. 2A, fig. S7, Movie S1) and recapitulates key growth patterns in ‘digital DiI’ simulations (Fig. 2B, Movie S2). In sum, the model describes the morphological transformation of the anterior neural tube to the characteristic form of the amniote forebrain.

**Figure 2:**
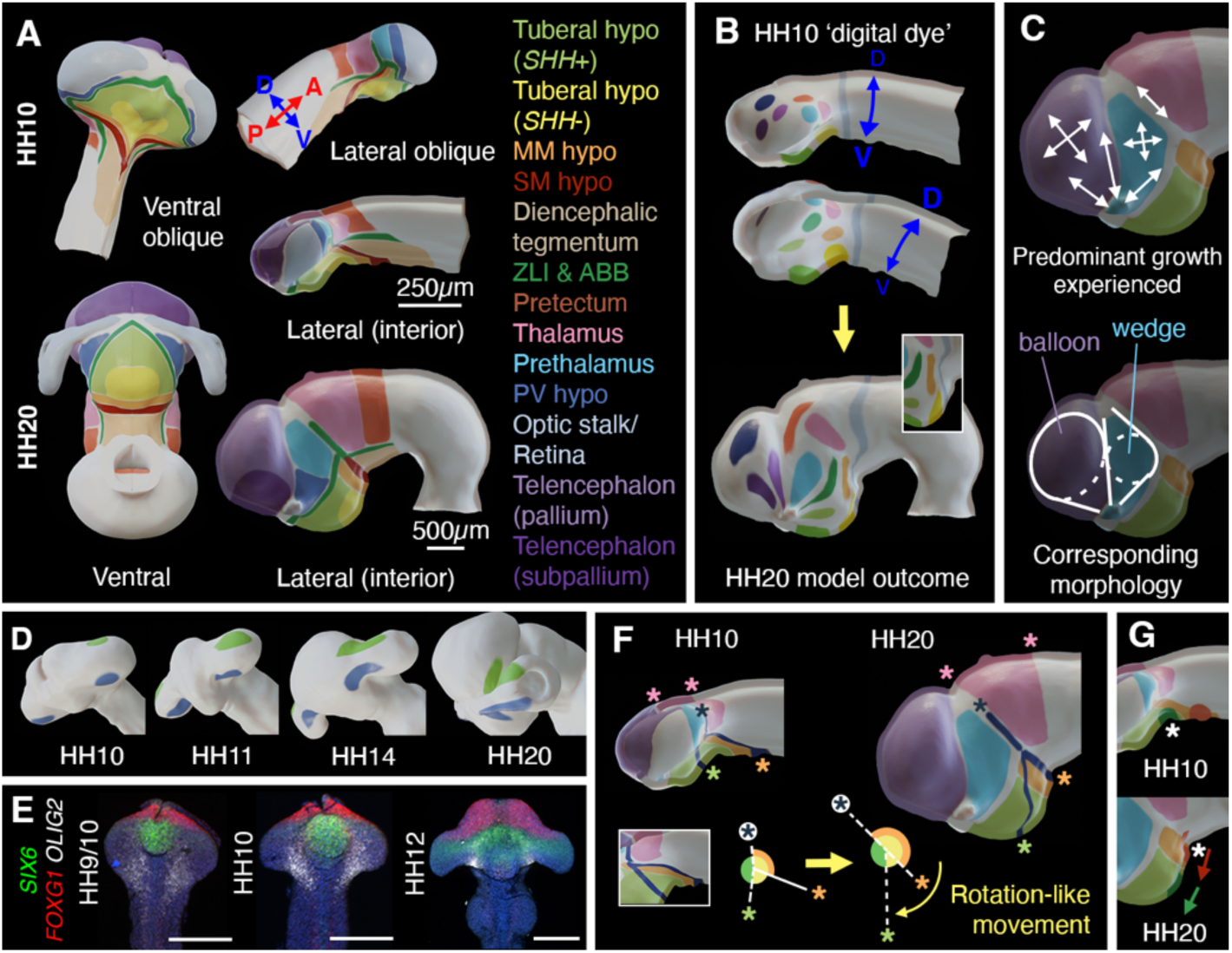
4D model of forebrain growth. **A)** HH10 fate map and forebrain regions at HH20. **B)** ‘Digital dye’ applied at HH10 (ventrolateral and dorsolateral views shown to aid visualisation) with resulting growth patterns. **C)** Simplified growth patterns (upper), and morphology resulting from isometric (circles) and directional (lines) growth (lower). **D)** ‘Digital dye’ spots progress to growth lines connecting eye and forebrain; oblique anterior-ventral views. **E)** HCR-stained neuroectoderms, ventral views. **F)** Model depicted as simplified fate map with dark lines and asterisks tracking position of thalamus and posterior hypothalamic cells relative to the ZLI. Lines intersect at a ‘hinge’-like point. **G)** Movement of ‘digital dye’ spots (green, red; arrows) in relation to HH10 ventral inflection point (upper, asterisk) and HH20 cephalic flexure (lower, asterisk).

Anterior to the ZLI, elongated growth lines converge towards the optic stalk, demonstrating that eye outgrowth is associated with strongly directional growth over a wide area of the forebrain (Fig. 2B, Movie S3 0:16-0:24). Up to HH20, anterior forebrain morphogenesis is dominated by the combination of this directional growth, and a more uniform expansion of the telencephalic pallium and the prethalamus (Fig. 2C, Movie S3 min 0:35-1.00). The telencephalon thus takes on a balloon-like shape, narrowing towards the optic stalk; the prethalamus and PV hypothalamus together become wedge-shaped, narrowing towards the dorsal midline and optic stalk (Fig. 2C); the optic vesicle/cup/stalk opening holds to a progressively smaller corner of the anterior forebrain (fig. S8A-D, Movie S3 1:03-1:20). Telencephalic and basal hypothalamic lineages are segregated from an early stage by the intervening prethalamus, PV hypothalamus, and eye field (Movie S3 1:22-1:47; compare growth lines in Fig. 2B to regions in Fig 2A). Growth lines connect regions sharing either *FOXG1* (telencephalon and nasal retina) or *FOXD1* (hypothalamus and temporal retina) expression (Fig. 2D-E, Movie S3 1:48-2:30, Fig. 1K, fig. S8E).

More posteriorly, the thalamus expands, contributing to the dorsal curvature of the neural tube (Fig. 2F, pink asterisks; Movie S4 0:08-0:18). Cells ventral to the ABB are displaced anteriorly by extension of the ventral midline, bringing them towards the telencephalon and creating V-shaped growth lines (Fig. 2F, green/orange asterisks; Movie S4 0:35-1:30. This movement resembles a rotation of ventral regions anteriorly, with a ‘hinge’ point at the intersection of the ABB and ZLI (Fig. 2F; Movie S4 1:34-1:48). The epithelial folds in the HH10 ventral midline contribute to the tuberal hypothalamic midline, and the ventral inflection point of the HH10 neural tube does not correspond to the HH20 cephalic flexure (Fig. 2G, asterisks; Movie S4 1:51-end), as is sometimes assumed.

### Repositioning of the dorsoventral and anteroposterior axes

Proponents of the prosomere model argue that higher proliferation in the dorsal neural tube (*17*, *19*) distorts block-shaped proto-segments into wedge-shaped prosomeres, causing the neural tube to pivot around the cephalic flexure (Fig. 3A) (*20–23*). This provides the rationale for the position of the A-P and D-V axes (Fig. 3B), and the positioning of the hypothalamus ventral to the telencephalon and anterior to the prethalamus (Fig. 3C). Our model does not support this claim: when purported prosomere boundaries (*16*) are drawn onto our model at HH10 and HH20, their positions are not as predicted when viewed at HH20 and HH10 (Fig. 3D). Instead, the bulk of the hypothalamus arises posterior to the telencephalon and eye, and ventral to the PV hypothalamus, prethalamus and ZLI (Fig. 3D-E).

**Figure 3:**
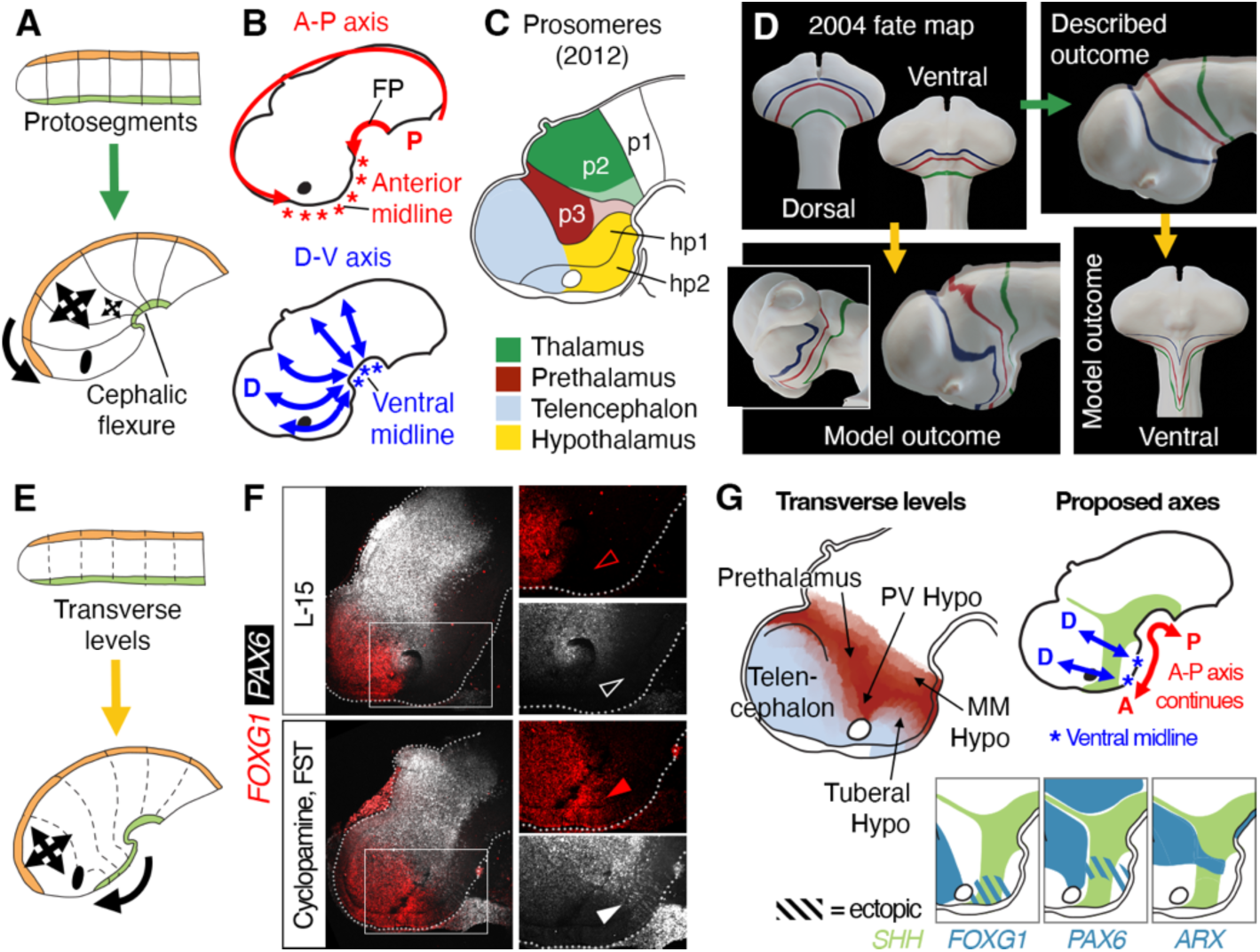
Repositioning of the dorsoventral and anteroposterior axes. **A)** Growth model underlying prosomere model. Two-headed arrows - differential growth, dorsal versus ventral regions. **B)** Prosomere model axes. **C)** 2012 prosomere model segments. P1-3 - diencephalic prosomeres 1-3, hp1-2 - hypothalamo-telencephalic prosomeres 1-2 (*4*, *6*). **D)** 2004 prosomere model boundaries at HH10 as per (*16*) (top left), with expected outcome (green arrow) at HH20 (top right). 4D model outcome (gold arrows) of the same prosomere lines at HH20 (bottom left) and HH10 (bottom right). **E)** Revised growth model. Two-headed arrows - telencephalic expansion. **F)** Control (L-15) and dorsalised (Cyclopamine, FST) embryos 24 hours after treatment at HH10, showing absent (open arrowheads) and ectopic (filled arrowheads) hypothalamic expression of *FOXG1* and *PAX6*. **G)** Revised A-P/D-V axes (top), ectopic expression of dorsal genes following dorsalisation (bottom).

The claimed shared segmental identity between the telencephalon and hypothalamus (Fig. 3C) implies that fate conversion should be possible only between these identities. Experimental dorsalisation of the hypothalamus using cyclopamine, or a combination of cyclopamine and follistatin (*7*), resulted in a small *SHH^+ve^* basal hypothalamus and decreased expression of *NKX2-1* (fig. S9). The telencephalic marker *FOXG1* was ectopically expressed, but only in part of the anterior tuberal hypothalamus; instead the prethalamic/PV hypothalamic marker *PAX6* was expressed ectopically in more posterior parts of the hypothalamus; *ARX* and *OLIG2* remained confined to the prethalamus, posterior tuberal and MM hypothalamus (Fig. 3F, fig. S9).

These shared competences are consistent with our fate map (Movie S1), as cells originating close to one another are exposed to similar patterning signals. We suggest that the anterior-most tuberal hypothalamus is topologically ventral to the telencephalon, in partial agreement with the prosomere model, but that the remainder of the tuberal region is ventral to the prethalamus and PV hypothalamus (Fig. 3G). The D-V axis within the hypothalamus is therefore rotated compared to the prosomere model, and the ventral midline - hence the A-P axis - extends into the anterior tuberal hypothalamus (Fig. 3B,G).

### A transverse boundary splits the hypothalamus and becomes distorted by differential growth

The ZLI is a lineage-restricted dorsal diencephalic signalling centre that divides the vertebrate nervous system into anterior *SIX/FEZ^+ve^* and posterior *IRX^+ve^* regions (*24–26*). Intriguingly, several genes in the HH20 posterior hypothalamus run approximately parallel to the ZLI and our proposed D-V axis. These include *PITX2* and *LMX1B* in the SM hypothalamus, *EMX2*, *OLIG2*, *WNT8B, BARHL2, EPHA7,* and *FOXD1* in the MM hypothalamus, and *LHX6*, *ARX* and *DLX1/2* in the prethalamic-like tuberomammillary terminal (TT) (Fig. 4A-E; Fig. 1E,K; fig. S1E; fig. S10A-F) (*7*). *PITX2* also marks a distinct anterior domain of the *FOXA1/LMX1B/SHH^+ve^* FP that abuts the *ARX^+ve^*FP at a flexure point (Fig. 4C, arrowhead; fig. S10B,G). The *PITX2^+ve^*SM is flanked by, and partly overlaps with, *WNT8B*, *SIM1*, *EPHA7* and *DBX1;* these genes converge at the ZLI and FP, *WNT8B* resolving from a wider uniform domain at earlier stages (Fig. 4D; fig. S10C-E,H-J).

**Figure 4:**
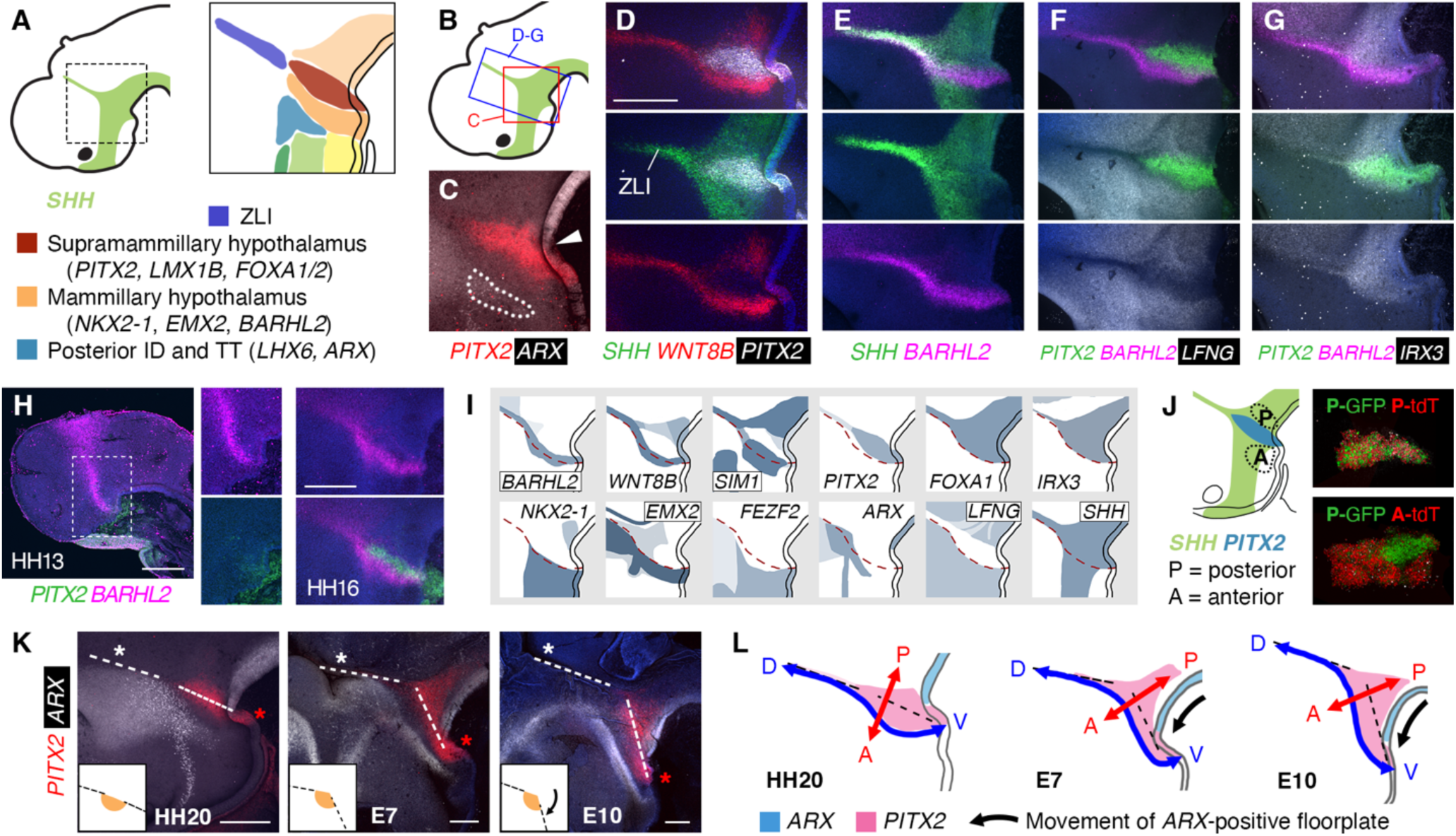
Dorsoventral domains in the embryonic posterior hypothalamus. **A)** Schematic of ZLI and posterior hypothalamic regions. **B)** Location of regions shown in **C-G). C-H)** HCR analysis of ZLI and posterior hypothalamic genes. **C**) Arrowhead - flexure where *ARX+ve* FP abuts *PITX2+ve* FP. Dotted outline - *ARX* in TT. **I)** Selected gene expression patterns, posterior hypothalamus and surroundings. Dotted line - edge of *LFNG-ve/IRX3+ve* area. **J)** Regions dissected for cell-mixing experiments indicated by dotted lines. Posterior GFP cells mixed with posterior tdTomato cells do not resolve (n= 7/7); posterior GFP mixed with anterior tdTomato separate (n= 6/8). **K)** HCR, HH20-E10 hypothalamus and ZLI. Dotted lines through ZLI (white asterisks) and *PITX2* SM region meet at an angle, resembling a ‘hinge’ point (inserts). Red asterisks - *PITX2+ve* FP. **L)** Proposed continued anterior movement of the FP distorts the A-P and D-V axis. Scale bars - 250 μm.

Strikingly, *WNT8B* and *BARHL2* encompass the ZLI dorsally and extend continuously through the hypothalamus to the *PITX2*/*SHH^+ve^*ventral midline (Fig. 4D-E, fig. S10I-J). A slender, straight *LFNG^-ve^* domain encompasses the ZLI (*27*), *PITX2^+ve^* SM, and part of the hypothalamic *WNT8B/BARHL2^+ve^*domain (Fig. 4F, fig. S10J,K). The *PITX2^+ve^* SM is *IRX1/IRX3^+ve^* and *FEZF2^-ve^*, as in mouse (*8, 28*, *29*). *BARHL2* expression straddles the edge of *IRX3* in the MM and ZLI (a slight kink is less pronounced at earlier stages, prior to the onset of *PITX2* expression)(Fig. 4G-H fig. S10H,L-M). Considered alongside the posterior origin of these cells (Fig. 2A), this ‘posterior’ molecular profile places the SM hypothalamus posterior to the prethalamus and contiguous with the ZLI (selected expression patterns are summarised in Fig. 4I). Basal plate cells taken from anterior and posterior to the SM segregate when mixed (Fig. 4J), suggesting that lineage restriction may operate in this region, as shown for an *En1-Dbx1* FP boundary in mouse (*30*). Together, we propose that a topological boundary, and hence the local D-V axis, traces the edge of the ZLI and the *LFNG^-ve^/IRX3^+ve^* hypothalamus at HH20, when hypothalamic patterning is largely complete (Fig. 4I, dotted line) (*7*).

At later developmental stages (E7-E10) many genes including *PITX2*, *ARX*, *WNT8B, LFNG, EPHA7, FOXA1, OTP, SIM1* and *FOXA2* bear similar topological relations to one another as at HH20 (Fig. 4K, fig. S11A-E). However, similar to the anterior movement of ventral cells at earlier stages (Fig. 2), the FP has shifted anteriorly relative to the ZLI. This creates an obtuse angle between the MM/SM domains and the ZLI, recalling the ‘hinge point’ in our growth model (Fig. 4K, insets; Fig. 2F). The D-V/transverse alignment of these regions becomes distorted and the MM/SM areas start to take on an A-P oriented/longitudinal appearance (Fig. 4L), the anterior edge of *WNT8B* aligning with a characteristic morphological fold, the hypothalamic periventricular organ (fig. S11F) (*31*, *32*). The topology of the embryonic forebrain thus becomes obscured by differential growth.

### Evidence for conserved growth patterns

The prosomere model partitions the hypothalamus along intersecting transverse (interprosomeric; D-V) and longitudinal (A-P) boundaries (Fig. 5A), proposed mainly on the basis of mouse gene expression patterns between E11-E19 (Fig. 5B; see fig. S12 for comparative hypothalamic terminology and selected regional marker genes)(*4*, *6–8*, *10*, *23*, *29*). The prosomeric A-P axis, not the D-V axis, follows the posterior hypothalamic expression domains of genes such as *Sim1* and *Otp*, which in mouse are highly curved and jut out almost perpendicularly to the ZLI (Fig. 5B). Given that the chicken D-V axis becomes distorted, we predicted that analysis of earlier mouse embryos might uncover a similar deformation.

**Figure 5:**
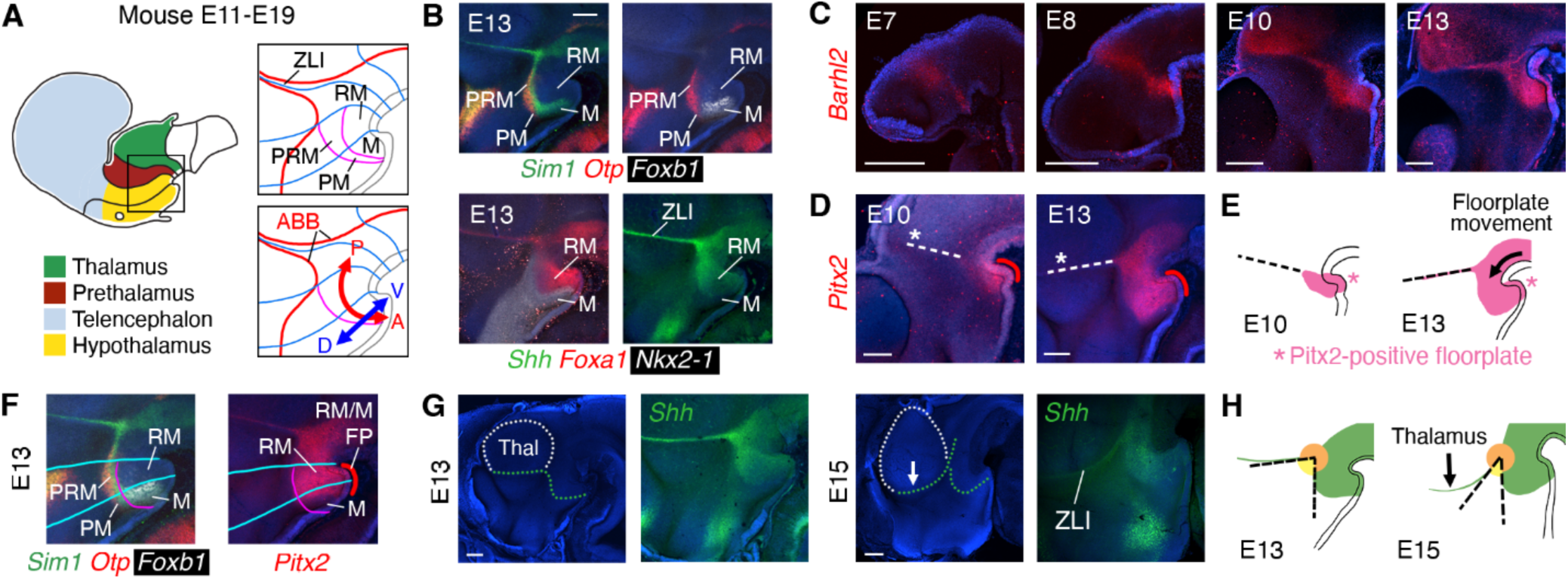
Growth patterns may distort the mouse forebrain axes. **A)** Mouse prosomere regions, boundaries (blue - transverse, magenta - longitudinal) and axes. **B)** HCR-stained E13 mouse (boxed region in 5A). **C-D)** HCR-stained E7-E13 mouse forebrains. Dotted line in (D) indicates position of ZLI, red line indicates *Pitx2*^+ve^ FP. **E)** Inferred movement of FP. **F)** Prosomere boundaries as per (A) overlaid on mouse E13 HCR. Red line - *PITX2*^+ve^ RM/M FP. **G)** DAPI- and (same samples) HCR-stained E13 and E15 mouse forebrains. Expansion of the thalamus (white dotted outline) displaces the *Shh*^+ve^ ZLI. Green dotted lines - anterior edge of *Shh* domain in ZLI and posterior hypothalamus. **H)** The angle between the ZLI and posterior hypothalamic *Shh* expression domain becomes more acute between E13 and E15. PM - perimamillary, PRM - periretromamillary, M - mamillary, RM - retromamillary. Thal – Thalamus. Scale bars - 250 μm.

At E7-E8 a transverse band of *Barhl2* indeed connects prospective thalamus with ventral posterior hypothalamus, an angle developing between the ZLI and hypothalamus by E10 (Fig. 5C). *Pitx2* becomes evident in the retromamillary (RM) and mamillary (M) hypothalamus (note prosomere terminology, fig. S12) from E10 (Fig. 5D), overlapping extensively with *Barhl2*. By E13 the *Pitx2^+ve^* FP has shifted anteriorly in relation to the ZLI (Fig. 5D-E), revealing that the *Pitx2^+ve^* FP underlies the RM and M hypothalamus (Fig. 5F). Simultaneously, thalamic expansion appears to thrust the dorsal ZLI forwards (Fig. 5G-H). Therefore, similar growth patterns to those revealed in chicken likely distort the mouse D-V axis even by E11, so that D-V oriented ventral regions (at E9-10) present as A-P oriented.

In parallel, we attempted to map prosomere model boundaries onto the embryonic chicken brain by examining defining prosomere regional markers (fig. S12C). This proved challenging: contrary to published accounts (*22*, *33*) the prosomere M domain could not be identified in the HH20 chicken, as transverse (using *SHH/NKX2-1/FOXA1*) and longitudinal (using *OTP/PITX2*) boundaries do not intersect as expected (Fig. 6A-B; fig. S12C) (*4*, *6*, *23*, *29*).

**Figure 6:**
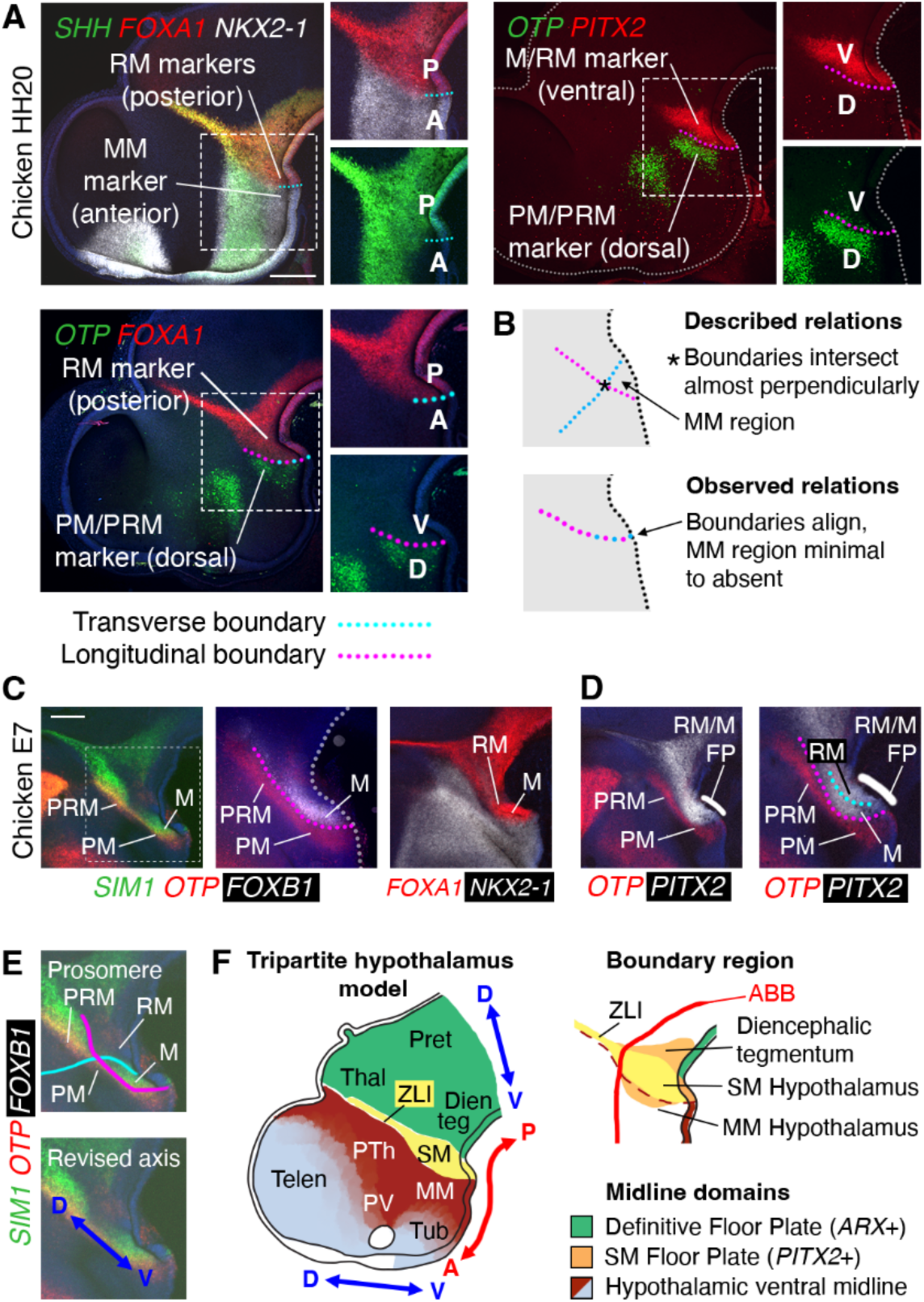
Chicken prosomere boundaries and a new model of forebrain organisation. **A)** HCR-stained HH20 chicken forebrain, prosomere regional markers and boundary positions. **B)** Described and observed (HH20 chicken) relations of prosomere boundaries. **C)** Identification of the chicken M domain and PRM-PM boundary at E7 (HCR). Dotted rectangle - area shown in middle panel (different sample). **D)** Estimation of the M-RM boundary using FP *PITX2* expression, E7 HCR. **E)** Result of mapping prosomere boundaries onto E7 chicken (upper), and our proposed D-V axis (lower). **F)** Revised forebrain model. Colours indicate transverse levels (left). Dotted line (upper right) - edge of *LFNG*^-ve^/*IRX3*^+ve^ domain. PM - perimamillary, PRM - periretromamillary, M - mamillary, RM - retromamillary. Telen - Telencephalon, PV - paraventricular hypothalamus, PTh - Prethalamus, Thal - Thalamus, Pret - Pretectum. Hypothalamus: Tub - tuberal, MM - mammillary, SM - supramammillary. Scale bars - 250 μm.

By E7, a slim *SIM1*/*FOXB1^+ve^* M region could be identified, but was orientated very differently to that in the E13 mouse, and overlapped with *FOXA1* (Fig. 6C). The RM-M transverse boundary, estimated based on the positions of *FOXB1* and FP *PITX2* (Fig. 6D, cyan line), ran almost parallel to the longitudinal M-perimamillary boundary (Fig. 6D, magenta line) rather than perpendicular. According to our axes (Fig. 4L), these two boundaries pass obliquely through the region that we designate mammillary (Fig. 6E, compare Fig. 5F) (*7*). The straighter edges of the *SIM1/OTP/BARHL2/PITX2* expression domains in chicken more closely resemble those in Xenopus, zebrafish and lamprey (*22*, *34–37*), suggesting that the curved morphology of the posterior hypothalamus in mouse is derived. We conclude that the prosomere model misinterprets the neural tube A-P and D-V axes due to an over-reliance on gene expression patterns from a single species at late developmental stages, combined with a flawed growth model arising from a lack of fate mapping information for ventral regions. Instead we propose a ‘tripartite hypothalamus’ model in which the hypothalamus sits ventral to three regions – the telencephalon, prethalamus/PV hypothalamus, and the ZLI (Fig. 6F). The *LFNG*^-ve^ ZLI and posterior hypothalamus form a transverse unit between the anterior and posterior forebrain, and the topological relations of forebrain regions reflect their early spatial origins - and therefore the environment seen by their progenitor cells.

## Discussion

We have completed the anterior neural tube fate map for HH10 chicken embryos and described the morphogenetic movements shaping the amniote forebrain. Evidence from fate mapping, multiplex HCR analysis, fate conversion experiments and morphology led us to reinterpret the forebrain D-V axis, with direct implications for the topological position of the hypothalamus, a source of major confusion due to its complexity and relative scientific neglect.

The anterior movement of ventral midline cells likely represents late gastrulation movements, driving the accumulation of tissue into epithelial folds (at HH10) and feeding cells into the basal hypothalamus. The ‘wedge’-shaped region encompassing the prethalamus and PV hypothalamus resembles the dorsal portion of the ‘D1’ neuromere of a previous study (*38*), and may provide the early morphogenetic basis for the (larger) optic recess region defined in zebrafish (*39*). Our results provide important context for understanding eye morphogenesis, a tissue that is very challenging to fate map owing to the rapid and extreme tissue deformations involved. Notably, our results are consistent with cell movements described in the zebrafish ventral midline (*40*, *41*), diencephalon (*25*), and eye (*42–45*), therefore helping to translate these findings into higher vertebrates.

We propose that the anterior tuberal hypothalamus is situated ventral to the telencephalon, and the posterior tuberal and anterior mammillary hypothalamus are ventral to the PV hypothalamus and prethalamus (Fig. 5K). The close relationship of much of the hypothalamus to prethalamus is well supported by both genetic and biochemical analysis (*7, 8, 10*, *11*, *46*). We identify a major forebrain boundary cutting through the MM hypothalamus and the ZLI, placing the SM hypothalamus (and part of the MM region) posterior to the prethalamus and ventral to the ZLI. We speculate that the SM hypothalamus and flanking *WNT8B*-positive regions represent an extensive ventral ‘boundary region’ (Fig. 5K), where the juxtaposition of anterior and posterior forebrain - plus underlying tissues such as notochord and prechordal mesoderm - provided an evolutionary substrate for the generation of high signalling and cell type complexity. Cells originating here contribute widely to the posterior hypothalamus and beyond, with *PITX2/LMX1B^+ve^*progenitors in mouse contributing to the SM, subthalamic and ventral premammillary nuclei (*8, 28*, *30*, *47*, *48*).

The SM FP is distinguished from ‘definitive’ (*ARX^+ve^*) FP by *PITX2* (Fig. 5K), likely conserved to mouse (*30*) and human (*49*). The rest of the hypothalamic midline lacks FP markers but arises from FP-like (HypFP) progenitors (*12*, *50*), its uniqueness accounted for by the distinct evolutionarily origin of the anterior forebrain (*51–54*), and mechanistically by the distinct patterning influence of the prechordal mesoderm (*55–57*). Our model incorporates elements from both columnar and prosomere models and, critically, is informed by evidence from early developmental stages, minimising the confounding effects of neuronal migration.

Although we conducted our fate mapping through patterning stages, when cell fate is initially plastic, the many similarities in the shapes of DiI growth lines and gene expression domains demonstrate the close coupling of growth and fate. Growth patterns help us to understand the epigenetic history and environmental exposures of cells during a crucial period of specification. Our results therefore offer new insights into forebrain morphogenesis and organisation and should help to inform efforts to direct pluripotent stem cell differentiation towards specific forebrain identities.

## Acknowledgments

The authors thank Alex Fletcher for comments on the manuscript, and Nick Van Hateren and Darren Robinson for help with imaging and Blender. Imaging work was performed at the Wolfson Light Microscopy Facility using the Zeiss Light sheet (BBSRC BB/MO12522/1) and Nikon spinning disc (BBSRC2020 Alert B/V019368/1) microscopes.

## Author contributions

Conceptualization: DWK, SB, MP, EP. Methodology: EM, KC, CF, MP, EP. Investigation: EM, KC, MP, EP. Visualization: EM, KC, CF, MP, EP. Formal analysis: EM, MP, EP. Validation: EM, MP, EP. Data curation: EM, MP, EP. Project administration: MP. Writing – original draft: EP. Writing – review: EM, MP, EP. Writing – editing: EM, KC, CF, DWK, SB, MP, EP.

## Data and materials availability

All data are available in the main text or the supplementary materials.

## Supplementary material

## Materials and Methods

### Embryo sourcing and incubation

Fertilised Bovan Brown eggs (Henry Stewart & Co., Norfolk, UK), and transgenic *Chameleon* (cytbow) (*15*), Cytoplasmic GFP (*58*), and Flamingo (tdTomato) (*59*) eggs (The Roslin Institute, Edinburgh, UK) were incubated in a humidified incubator at 37°C until the desired stage, according to (*60*). Time-mated pregnant wildtype C57B1/6 mice were sourced from Envigo RMS (UK) Limited, Shaws Farm, Bicester. All studies and procedures were conducted according to the UK Animals (Scientific Procedures) Act 1986/EU Directive 2010/63/EU, and were approved by the University of Sheffield Local Ethical Review committee. Ethical approval or Home Office licensing was not required for egg work, as eggs were not incubated beyond E10. Named Animal Care and Welfare Officers (NACWOs) had oversight of all incubated eggs.

### Immunolabelling

Embryos were fixed in 4% PFA for 2 hours at 4°C and washed in PBS. Samples were incubated in block solution (1% TritonX, 10% heat-inactivated goat serum (HINGS) in PBS) overnight, then primary antibody solution (Rabbit anti-laminin α (LAMA), Sigma L9393 at 1:1000 in block solution) overnight. Embryos were washed twice in PBS at room temperature, and transferred to 10% HINGS in PBS and left overnight. Alexa 488-conjugated anti-rabbit IgG secondary antibody (1:500, Jackson ImmunoResearch) and phalloidin-TRITC (1:100, 1% TritonX, 1% HINGS in PBS) were applied at 4°C overnight, washed the following day in PBS at room temperature, then washed overnight in PBS with 0.1% TritonX. All steps were performed at 4°C except where stated.

### DiI injections

Eggs were windowed and Coomassie Blue (0.5 μl/ml in L-15 - Fisher Scientific, Cat No 11580396) was injected under the embryo to aid visualisation. The forebrain inner neuroectoderm was accessed via a dorsal midsagittal incision. CellTrackerTM CM-DiI Dye (Invitrogen, Cat No. C7000) in ethanol (50 μg per 30 μl), or Vybrant™ DiO cell-labeling solution (Invitrogen, Cat No. V22886) was loaded into a fine glass needle and injected into embryos by hand using a Parker Picospritzer II (10-20 msec pulses, 15 psi). Dye location was noted and imaged using a Leica MZ16F stereomicroscope. Embryos were incubated for 48h or as stated, fixed in 4% buffered paraformaldehyde, hemisected and reimaged prior to processing for HCR. Some DiI/DiO signal is lost during the HCR procedure, therefore dye and gene expression patterns are shown separately for some examples.

### HCR analysis

Embryos were fixed overnight in 4% paraformaldehyde at 4°C and stored in methanol for a minimum of one night before rehydrating in PBS. HCR v3.0 (*61*) was performed on intact embryos according to the manufacturer’s RNA-FISH protocol for whole-mount chicken and mouse embryos (as shown on Molecular Instruments website), using reagents obtained from Molecular Instruments, Inc. (Los Angeles, CA, USA). HCR probes were designed by the manufacturer based on the following accession numbers: **Chick**: *ARX* (XM_025146483.1), *BARHL2* (XM_015290665.4), *EMX2* (XM_025152058.1 and XM_025152057.), *EPHA7* (NM_205083.1), *FEZF2* (XM_414411.5), *FOXA1* (XM_004941922.3), *FOXA2* (NM_204770.1), *FOXD1* (NM_205192.3), *FOXG1* (NM_205193.1), *IRX3* (XM_015292372.4), *LFNG* (NM_204948.1), *LHX6* (XM_015279838.2), *LMX1B* (NM_205358.1), *NKX2-1* (NM_204616.1), *NKX2-2* (XM_015283379.2). *OLIG2* (NM_001031526.1 with extra sequence from NC_006088.5|:106522977-106525323 #256 chromosome 1, GRCg6a), *PAX6* (NM_205066.1), *PITX2* (NM_205010.1 with additional 5’ exons from XM_025149516.1/ XM_025149515.1), *SHH* (NM_204821.1), *SIM1* (XM_004940357.3), *SIX6* (NM_001389365.1), *WNT8B* (XM_025151998.1). **Mouse**: *Barhl2* (NM_001005477.1), *Foxa1* (NM_008259.4), *Foxb1* (NM_022378.3), *Nkx2.1* (NM_009385.4), *Otp* (NM_011021.5), *Pitx2* (NM_001287048.1), *Shh* (NM_009170), *Sim1* (NM_011376.3).

Proteinase K treatment (10 µg/ml) was performed for 2-30 min and embryos were re-fixed for 20 minutes, washed in PBST and transferred to 5x SSCT on ice. Embryos were preincubated in hybridisation buffer for 30 mins, then hybridised overnight in 10 nM (1:100) probe solution in hybridisation buffer, both steps at 37°C. Samples were washed 4 x 15 minutes in wash buffer at 37°C, then 2 x 5 minutes in 5x SSCT at room temperature. Embryos were equilibrated in amplification buffer for 5 minutes before adding the hairpins solution. Even and odd hairpins were melted (90 secs, 96°C) and cooled (30 minutes, room temperature) separately before mixing in amplification buffer (1:50). Samples were incubated in the dark overnight and washed in 5x SSCT (2 x 5 mins, 3 x 30 mins, 1 x 5 mins). Embryos were hemisected or the neuroectoderm isolated (see below) as necessary. Tissue was counterstained with DAPI prior to imaging (Cell Signaling Technology, Cat no. 4083, 1:1000).

### Cre recombination and neuroectoderm isolation

Chameleon (Cytbow) eggs were incubated until HH9, windowed, staged, and screened for the Cytbow transgene, using a handheld 365 nm black-light torch. 0.5-1.5 μl TAT-Cre recombinase (1500U) (Merck Life Science UK, SCR508) diluted 1:20 to 1:200 in L-15 was injected into the anterior neural tube using a pulled capillary needle. Eggs were sealed and incubated for a further 24-48 hours before fixing and staining for *NKX2-2/WNT8B* by HCR. Following HCR, forebrains were hemisected and the neuroectoderm isolated by microdissection aided by treatment with Dispase (1 mg/ml) for approximately 10 minutes to help separate tissue layers. Highly curved regions were nicked with a scalpel to enable flat mounting on a standard microscope slide.

### Electroporations

A Microelectrode holder (World precision Instruments, MEH6RFW) with a 0.25 mm silver wire was used as the cathode. A capillary needle containing 1-2 ug/ul pCAGGS-RFP with 0.1% Fast Green dye was threaded over the silver wire, and a mouth pipette attached to the side inlet valve to control the internal pressure. The anode, made from a tungsten needle, was positioned under the target cells and 2×3 pulses of 40V were applied using a TSS20 Ovodyne electroporator (Intracel), whilst simultaneously increasing the pressure in the needle to release the DNA mixture. Eggs were then re-sealed and incubated for a further 24-48 hours.

### Imaging and image processing

Images were taken on a Zeiss Apotome 2 (10x objective) with Axiovision software (Zeiss), Leica MZ16F using LASX software, Zeiss Lightsheet (5x, 10x objectives) with ZEN software (Zeiss), and Nikon W1 Spinning Disk confocal (4x, 10x, 20x objectives) with Nikon Elements software. For Apotome, Lightsheet and Spinning Disc imaging, embryos were mounted in 1% agarose, and z-stacks are presented as maximum intensity projections. Image processing was performed using Fiji (ImageJ, version 2.7.0/1.53t) and Adobe Photoshop 2022. Hemisected embryos are presented as the right side of the embryo, for ease of comparison. Gamma levels have been altered for clarity.

### 4-dimensional modelling of forebrain morphogenesis

Neuroectoderms were isolated (see above) from HH10, HH11, HH14 and HH20 chicken embryos, stained with DAPI, and imaged on a Zeiss Lightsheet confocal (fig. S6, step 1). Image stacks were converted to binary, denoised, and exported as a wavefront (.obj file) using FIJI software and the 3D viewer plugin (fig. S6, step 2). The .obj was imported into the open source animation software Blender (blender.org) and the smooth tool (in sculpting mode) was used to remove the artefactual contour lines. The objects were remeshed to reduce file size, and these meshes were used as morphological templates to build the model around (fig. S6, step 3).

A simple mesh was built around the HH10 template by first adding a single plane and enabling snapping to the surface of the template (using Face Project, with ‘Project individual elements’ enabled, as well as the Shrinkwrap modifier; Backface culling was enabled, as was ‘in front’ under Object Properties/Viewport Display). Geometry was added using the modelling tools, until the entire right hand side of the neuroectoderm was represented by a simple quad mesh. The mirror modifier was used to view the entire neuroectoderm to assist progress, but never actually applied to the mesh. Different faces of the mesh were given different colours, to assist subsequent stages by making them more easily identifiable. The simple quad mesh was then duplicated (ensuring the new object was not linked to the original) and snapped to the surface of the HH20 template mesh. The vertices were moved using the tweak tool, so that their positions at HH10 and HH20 roughly corresponded, according to the DiI/DiO results from the present study and the fate maps of Garcia-Lopez *et al*. (*16*) and Pombero *et al*. (*17*). In a similar manner, the meshes were copied and snapped to the HH11 and HH14 template meshes, and the vertices positioned to intermediate points (between the HH10 and HH20 meshes) (fig. S6, step 4). The meshes were periodically subdivided by applying the subdivision modifier (simple setting, fig. S6, step 5) and adjusting further, until the level of detail and precision, was deemed sufficient. All such changes to the mesh geometry were applied identically to all four meshes in order to enable joining later. Progress was monitored along the way by joining the meshes (usually only the HH10 and HH20 meshes) as shape keys (‘Join as shapes’ option); selected faces were then coloured according to HH10 position or HH20 region, and the outcome at the opposite timepoint was viewed to assess the fate map or ‘digital dye’ growth lines.

Once the vertex positioning was complete, the four meshes were duplicated to the insides of the template meshes and adjusted for optimal fit. Inner and outer meshes were joined and subdivided further, so that they closely followed the surfaces of the template meshes (fig. S6, step 6). The four meshes were then joined as shape keys to enable warping between the four stages (fig. S6, step 7). Fate map and ‘digital dye’ colours were applied to the surface by creating a UV map and using the texture painting options (fig. S6, step 8). A procedural material was assigned to generate a surface texture. Forebrain animations were created and exported from Blender, and annotated movies were compiled using Adobe Premier Pro software. Note that the modelling approach does not allow for any ‘cell mixing’ between adjacent parts of the mesh, hence ‘digital growth lines’ are slightly smaller than *in vivo*.

### Morphometric analysis

HH10-HH19 forebrains were stained by HCR for *FOXG1*, *OLIG2* and *WNT8B* and hemisected and imaged on a Nikon W1 Spinning Disc confocal. Morphometric measurements were performed on maximum projections of z-stacks using FIJI (ImageJ, version 2.7.0/1.53t). Distance measurements were obtained using the line tool and ‘measure’ function. First, line A (fig. S8B) was drawn between the trackable points X and Y: respectively, the midline at the ventral limit of FOXG1^+ve^, and a corner of the *OLIG2* expression domain in area 6 (fig. S8B-C; Fig. 1D,T). A perpendicular line was then positioned to intersect both line A and the posterior margin of the optic vesicle/stalk, which was clearly visible in hemisections as an edge (brightness/contrast levels were adjusted as needed). Line B was drawn between the posterior limit of A and the point of intersection of A and the perpendicular line. Results were exported to Excel using the Read_And_Write_Excel Plugin, the B:A ratio was calculated, and a logarithmic trend line was added.

### Dorsalisation of chicken embryos

Embryos were windowed at HH10-11 and the vitelline membrane torn above the head. Dorsalisation was induced by treatment with the SHH antagonist Cyclopamine (1 μg /μl; Sigma, dissolved in PBS with carrier 45% 2-hydroxypropyl-β-cyclodextrin, Sigma), and the TGF-ꞵ inhibitor Follistatin, recently shown to repress ventral identity (*7*) (15 μg/μl Recombinant Mouse Follistatin 288; 769-FS, Bio-Techne). 5μl of cyclopamine only or 5μl of Cyclopamine/FST (1:1) was pipetted on top of the embryo and the egg was re-sealed. Embryos were dissected and fixed 24-48 hours later and processed for analysis by HCR.

### Cell mixing analysis/ Fish ball

Transgenic Cytoplasmic GFP and Flamingo (tdTomato) chicks were dissected at HH16-17 into L-15 on ice. Neuroectoderm were isolated following Dispase (Sigma-Aldrich, Cat No. 4942086001) treatment and regions anterior (Td) and posterior (GFP) to the mammillary pouch were dissected respectively. The anterior and posterior explants were combined and incubated in Papain (2mg/ml) for 10 mins and dissociated into single cells by trituration using fire polished glass pipette with different capillary openings (large and small). Following dissociation, the enzyme was inactivated with 5 mls of L-15 and gently centrifuged at 600 RPM for 5 mins at 4°C. Cells were resuspended in 1 ml of explant medium (*62*) and counted using a haemocytometer. Hanging drops were formed from 10-15ul cell suspension containing 8000-10000 cells/ul, and incubated at 37°C for 24-48hrs to form pellets. Cell pellets were fixed, cryosectioned and imaged without further staining.

## Supplementary Figures

**Fig. S1.**
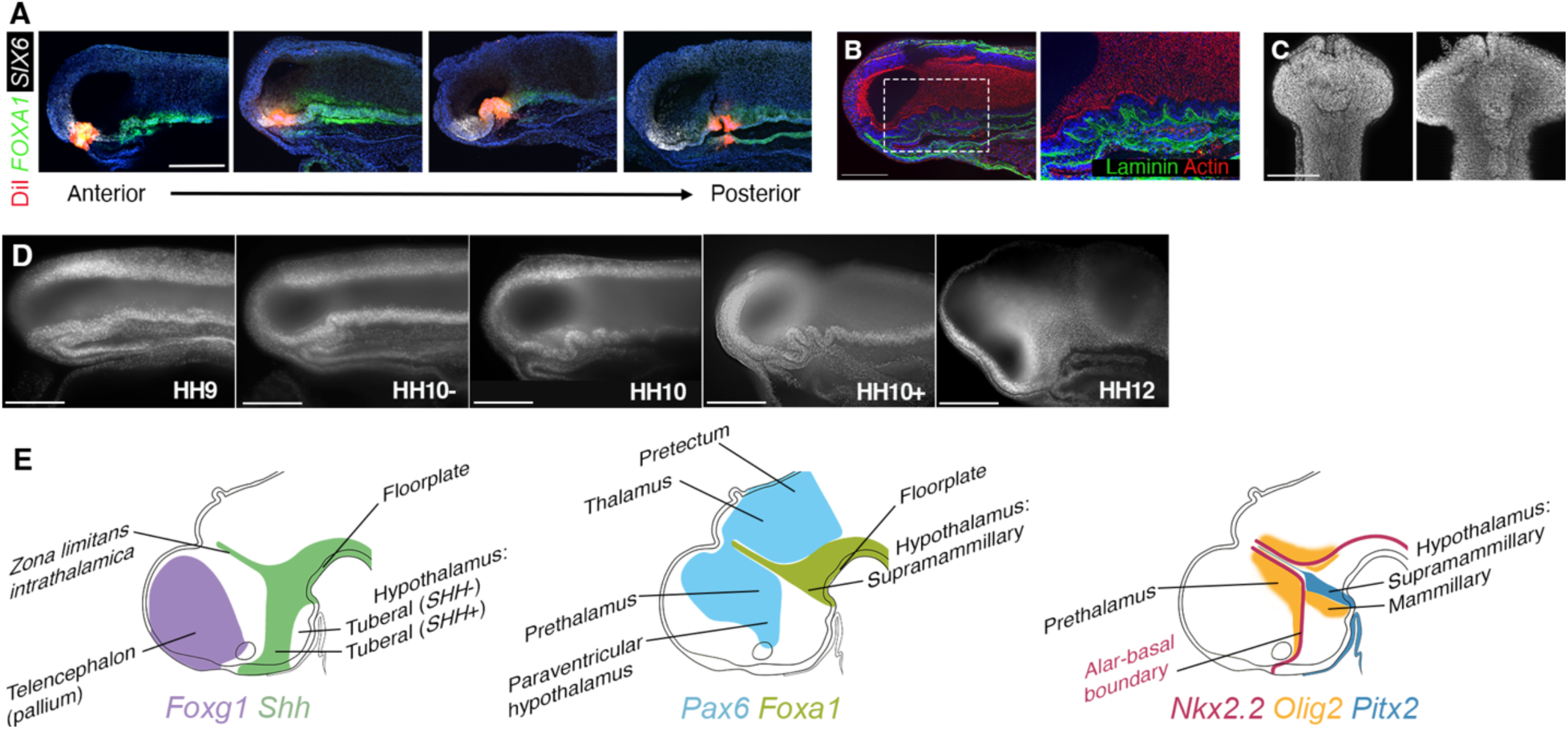
Morphological and genetic landmarks at HH10 and HH20. **(A)** Accurate targeting of DiI. Hemisected internal views of HH10 embryos analysed by HCR for expression of *FOXA1* (green) and *SIX6* (white) after DiI targeted to consecutive regions along the A-P ventral midline. **(B)** Hemisected internal view of HH10 embryo immunolabelled for laminin and stained with phalloidin to reveal Actin reveals midline epithelial folds underlain by basement membrane. Higher resolution of boxed area shown to right. **(C)** HH10 isolated neuroepithelia, DAPI-stained, ventral views. Midline epithelial folds increase in number and area from HH10- to HH10+. **(D)** Hemisected internal views of HH9-HH12 chicks, DAPI-labelled. Epithelial folds form, then resolve between these developmental stages. **(E)** Selected regional marker gene expression patterns.

**Fig. S2.**
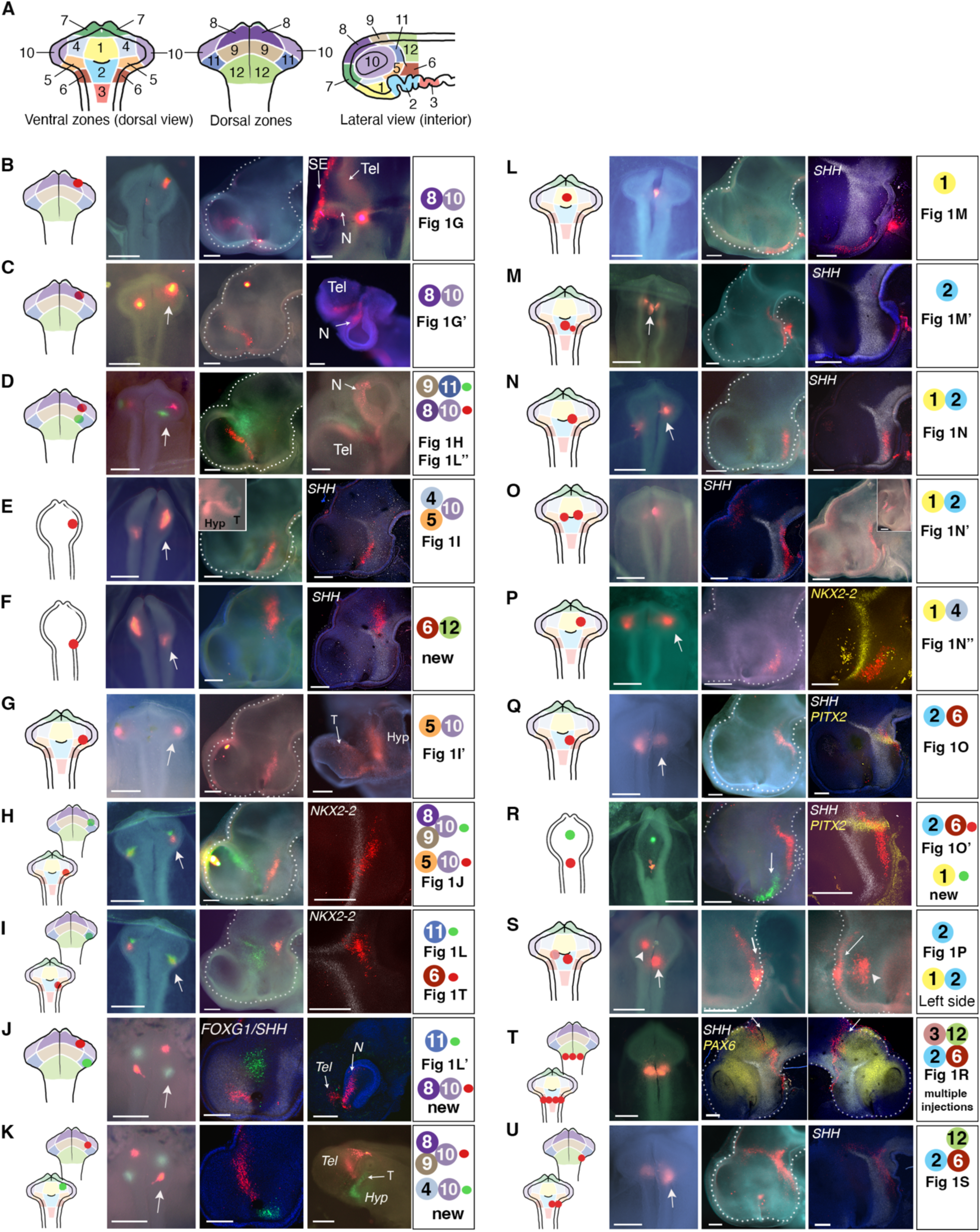
Regionally distinct patterns of anisotropic growth in the anterior neural tube. **A)** Different views of the HH10 forebrain, depicting the 12 regions targeted for injection. **(B-U)** From left to right: panels 1 show injection site(s); panels 2 show *in ovo* images of initial injection. White arrows point to injections shown at HH20 (panels 3, hemisected internal views). Panels 4 show hemisected internal views after HCR in situ to detect genes indicated (E-F,H-I,L-N,P-R,T-U), or show alternative views to aid with visualising particular growth lines (B-D,G,J-K,O(inset),S). Panels 5 indicate regions mapped and whether sample has been shown in Fig1, or is ‘new’. Where both sides of the embryo are shown (e.g. (E-F, H-I)), the left side is shown second (below); all injections and hemiviews are presented as right hand sides to assist comparison. Outer eye views in (D,G,J,K) have been flipped horizontally, to aid comparison with hemiviews. **(B-D)** DiI-labelled growth lines stretch from telencephalon to eye (panels 3) and into nasal eye (panels 4; anterior views); (D) also shows uniform expansion of DiO prethalamic territory. **(E, G, H)** DiI-labelled growth lines stretch in an AP direction towards eye, crossing the *NKX2-2^+ve^*ABB (from *SHH^+ve^* to *Shh^-ve^* territory) and entering the optic stalk (panels 3 and 4) to connect the hypothalamus and temporal eye (inset in (E) and panel 4 (G)). **(F)** Same embryo as in (E), DiI labelling in ventral thalamus. **(H)** DiO growth line stretches from the dorsal telencephalon to the eye. Together the DiI and DiO growth lines create a triangle that outlines the prethalamus and PVH. **(I)** Same embryo as in (H). Area 11 (DiO) labels the prethalamus, and Area 6 (DiI) injection forms a tricorn shape close to the ABB (marked by *NKX2-2*) and the ZLI (flanked by *NKX2-2*). **(J)** Note - 24 hour incubation period. Area 11 (DiO) labels prethalamus/PVH, Area 8/10 (DiI) labels *FOXG1*^+ve^ telencephalon and nasal eye. **(K)** Same embryo as in J). Area 8/9/10 boundary (DiI) stretches along the border of the *FOXG1^+ve^*telencephalon (compare with HCR in (J)). Area 4/10 injection stretches from anterior hypothalamus along the optic stalk, as shown in ventral view. **(L)** Area 1 midline injection labels anterior hypothalamic midline. **(M)** Area 2 midline injection labels tuberal *SHH^-ve^* midline. **(N, O)** Lateral Area1/2 injections label lateral tuberal hypothalamus. In (O), HH20 embryo has been cut unevenly so midline is missing. Inset shows left hand side of same embryo with growth lines from left and right side injections running parallel to each other along the lateral hypothalamus. **(P)** Area 1/4 injection in anterior hypothalamus. **(Q)** Area 2/6 injection bends around the floor plate to the tuberal/mammillary midline, crossing through the *PITX2^+ve^* region. **(R)** Estimated area 2/6 (DiI) and Area 1 (DiO) in a HH9 embryo. The dorsal regions have been removed from the embryo to visualise the ventral surface. As at HH10, Area 2 injections stretch along the tuberal ventral midline, becoming distorted posteriorly around the *PITX2* region. The anterior midline injection (DiO) labels the anterior *SHH^+ve^* hypothalamus, forming a V posteriorly (arrow). **(S)** Area 2 midline injection (arrow) and lateral Area 1/2 injection (arrowhead). Left and right sides of the HH20 shown, left side image has not been horizontally flipped. The midline injection forms a long V shape (arrows). **(T)** Multiple injections at different D-V positions at a single transverse level. Left and right sides of the HH20 shown, left side not horizontally flipped. Dorsal label runs along the D-V axis of the *PAX6*^+ve^ pretectum (right side) or pretectum-midbrain boundary (left side) (arrows). Ventral label is carried more anteriorly and enters the tuberal *SHH^-ve^* hypothalamus ventrally. **(U)** Large injection encompassing Areas 2, 6 and anterior 12. Dorsal label runs parallel to the ZLI in the thalamus with cells labeled at the dorsal tip of the ZLI spilling over into the prethalamus; ventral label extends at an angle towards the tuberal midline. The direction of predominant growth changes at the ABB. Hyp - hypothalamus, N - Nasal retina, SE - surface ectoderm, T - Temporal retina, Tel - Telencephalon. n> 225 injection sites in 112 embryos at HH10; N> 50 injection sites in 24 embryos at HH5-HH9.

**Fig. S3.**
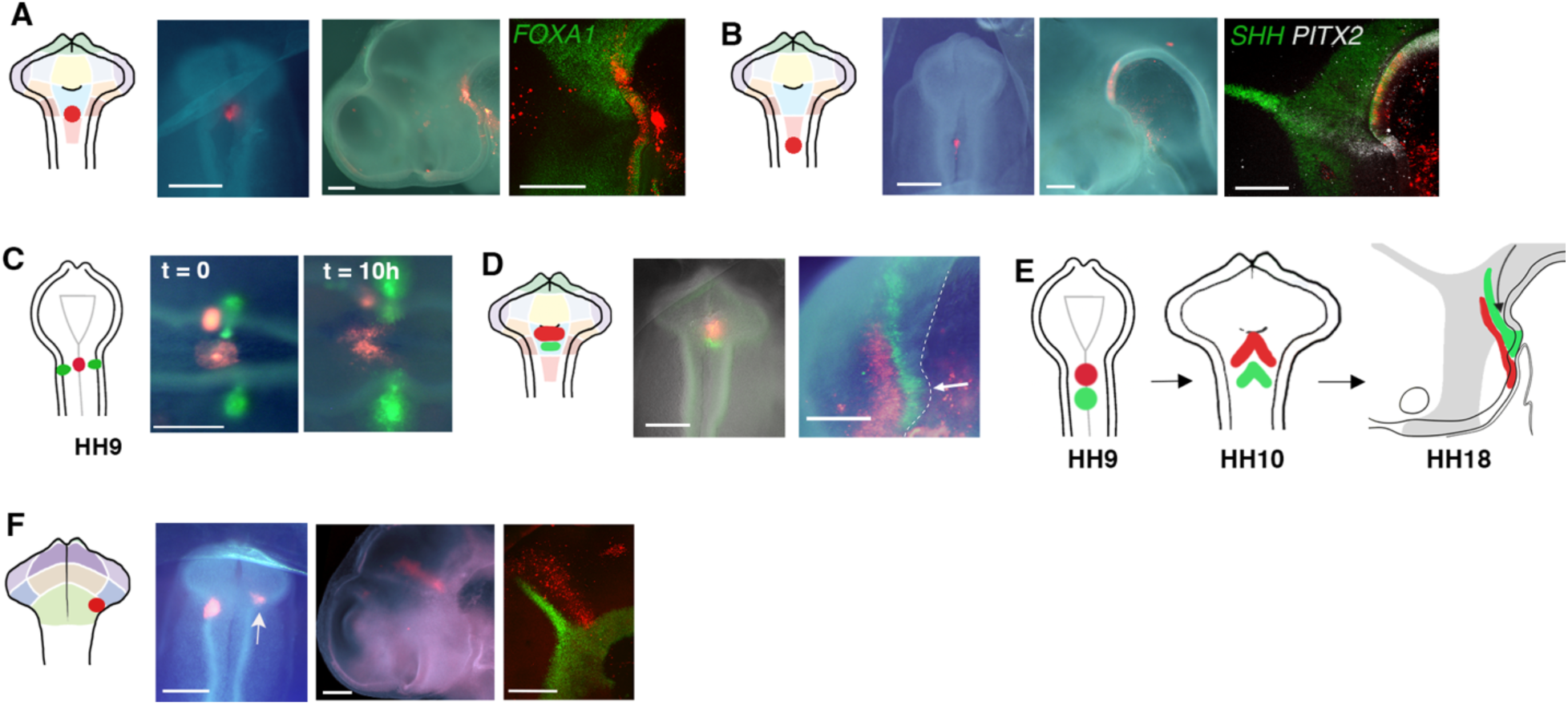
Midline and thalamic growth patterns (A, **B)** DiI injected embryos shown as schematic to indicate injection site, original injection at HH10, and resulting growth lines at HH18-20 (hemisected internal views). **(A)** Area 2/3 midline injection crosses from the *FOXA1^+ve^* FP to the hypothalamus. **(B)** Posterior area 3 midline injection contributes to the FP, posterior to *PITX2*. **(C)** Schematic shows dorsal view of HH9+ embryo after targeting DiI to ventral midline and DiO to more lateral regions at same transverse level. *In ovo* images shows at t = 0 and following 10 hours incubation (to ∼HH12). Midline DiI-labelled cells move anterior relative to lateral DiO-labelled cells, forming a V shape. Anterior to left. **(D)** Injection of DiO immediately posterior to DiI in Area 2 ventral midline. Resulting growth lines at HH18 form a nested V-shape. Arrow points to mammillary pouch. **(E)** Schematic of growth lines from HH9-18. **(F)** Area 12 injections immediately behind the ZLI expand along the D-V axis. Scale bars - 250 μm.

**Fig. S4.**
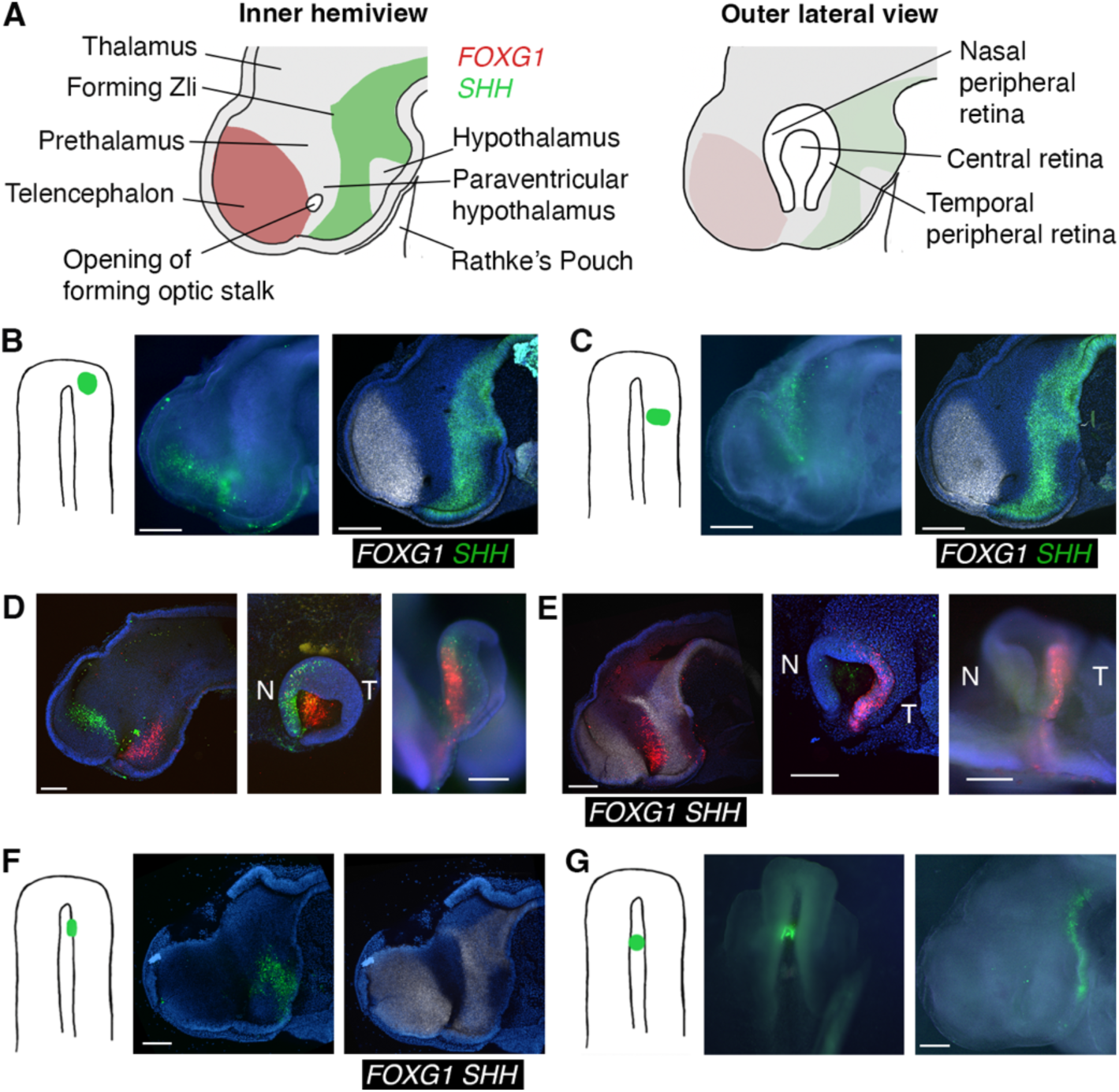
Growth lines resulting from HH8 injections. **(A)** Forebrain regions and marker genes at HH15. Inner hemiview (left) and outer, eye view (right). (**B-G)** Chicken embryos injected at HH8 and developed for 48 hours to approximately HH15-16. Schematics to the left show injection sites. HH15-16 embryos are shown before and after HCR processing. Note, DiO does not survive HCR processing. (**B)** Anterior injection forms elongated growth line in telencephalon. *FOXG1* (white) and *SHH* (green) shown on the same sample. **(C)** Left side of same embryo shown in (B), displayed as right side. A more posterior injection labels the prethalamus and PV hypothalamus together. **(D)** Double-labelled embryo shows growth lines from ventral and dorsal regions stretching into the eye. The isolated neuroepithelium (middle, right panels) shows telencephalic growth line connecting to the nasal retina, and hypothalamic growth line connecting to the central retina. **(E)** Growth line stretching between the PV hypothalamus and temporal eye. **(F)** A medial injection at HH8 labels the tuberal hypothalamus. Most of the growth line slopes anteroventrally, with a smaller dorsal part directed towards the eye. **(G)** A midline injection at HH8 becomes highly elongated along the A-P axis, from the tuberal hypothalamic midline to slightly lateral posterior regions.

**Fig. S5.**
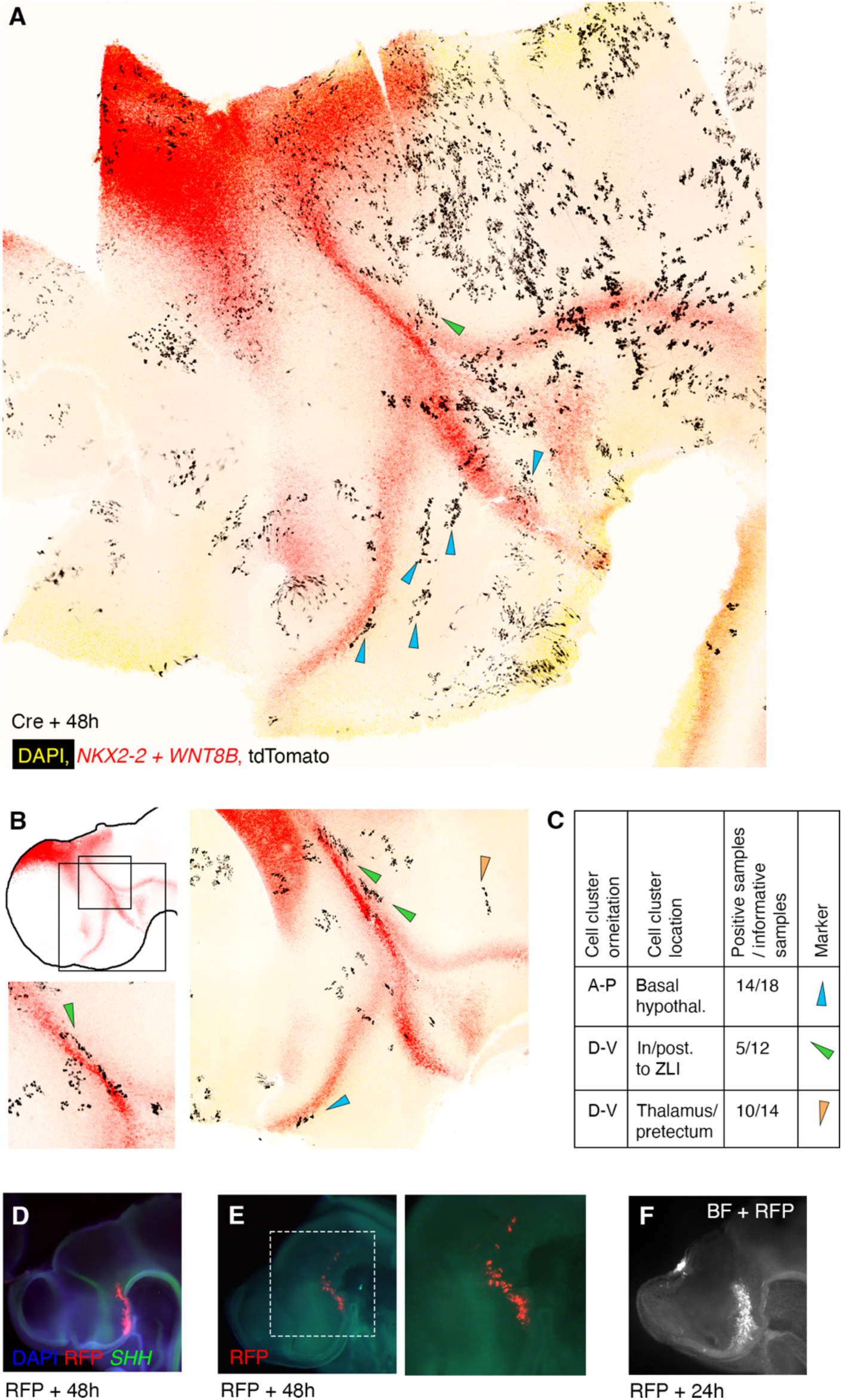
Cre-recombination lineage tracing and RFP electroporation confirms anisotropic growth patterns. **(A, B)** Flat mounted neurectoderm from Chameleon chick brains following injection of TAT-Cre recombinase into the lumen of the head at HH9. Following 48 hours of development (to ∼HH18), samples were processed for *NKX2.2* and *WNT8B* expression by HCR. Resulting tdTomato^+ve^ clones from Cytbow cassette recombination are shown in black. Blue arrowheads indicate clones that have extended along the A-P axis in the basal hypothalamus, green arrowheads indicates D-V expansion posterior to the ZLI, and orange arrowhead indicates D-V expansion in the thalamus/ pretectum. Boxes in schematic in (B) indicate areas shown below and to the right. **(C)** Quantification of the three patterns shown. Positive samples - the number of specimens containing cell group(s) showing this pattern. Informative samples - the number of specimens containing well-spaced cell groups in the specified region, that could therefore be assessed for their shape/orientation. **(D-F)** Targeted electroporation of an RFP construct at HH10 leads to distributions of labelled cells within the basal hypothalamus that resemble the shapes of growth lines seen with Dye labelling.

**Fig. S6.**
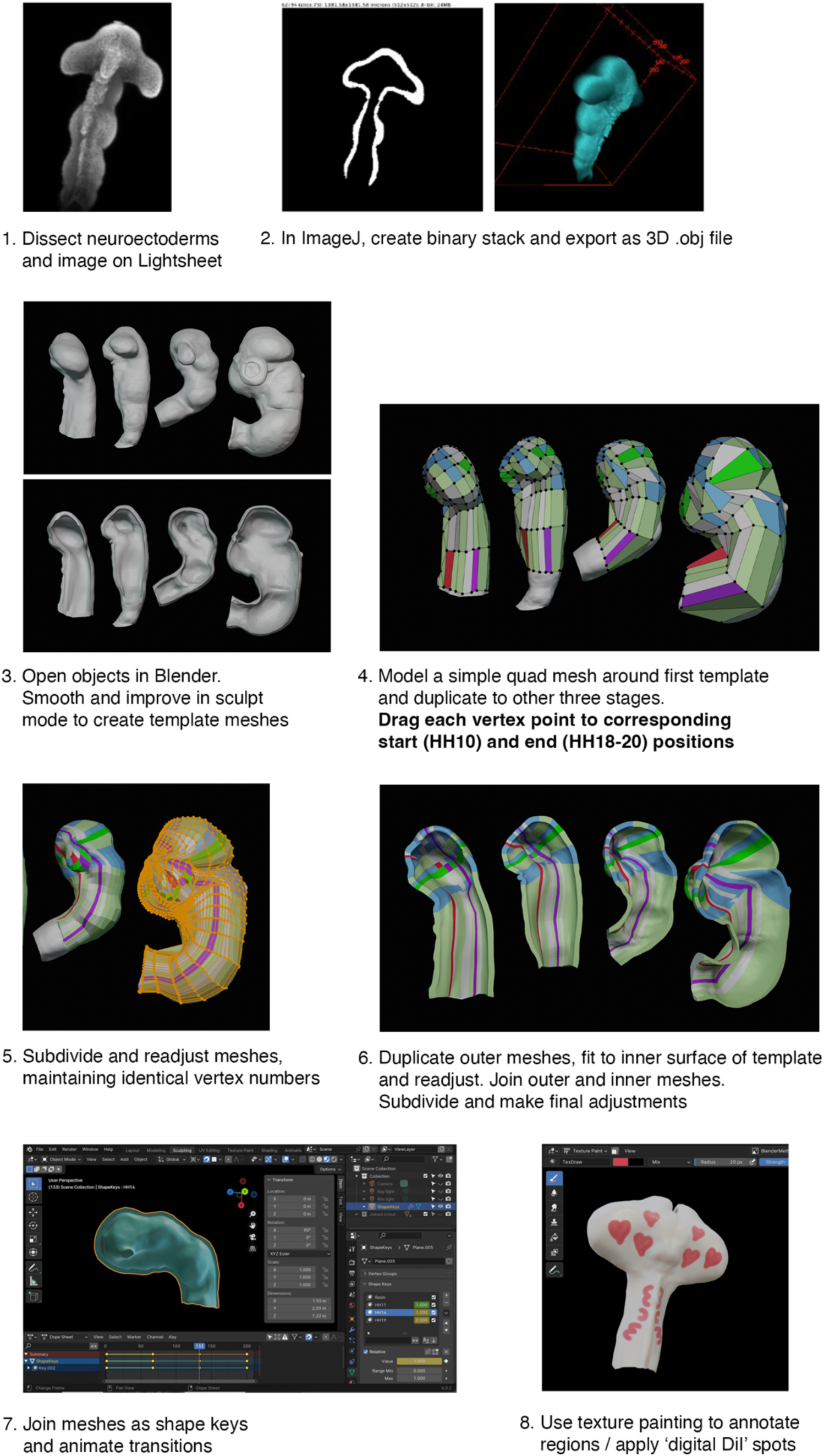
Building a 4D growth model. Stages involved in building the 4D forebrain model using Blender software (see methods for details). Scale bars - 250 μm.

**Fig. S7.**
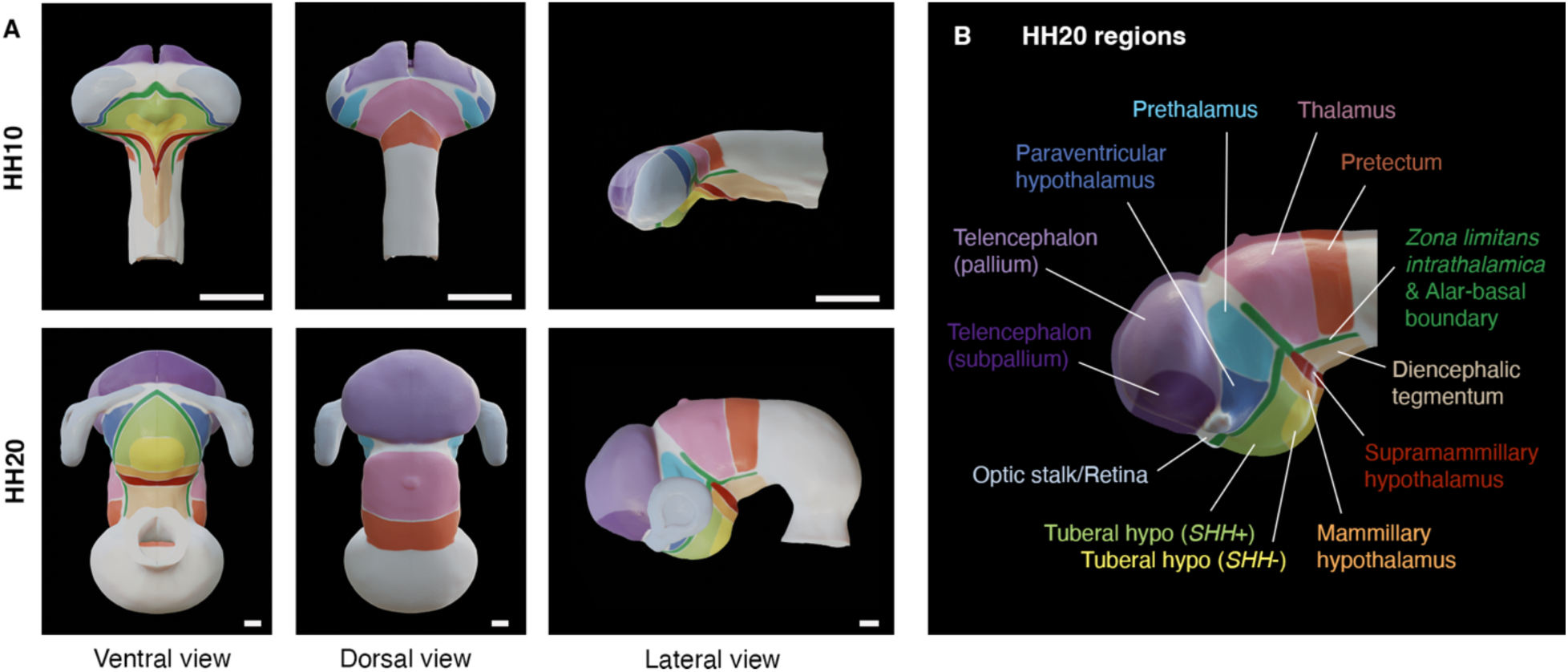
HH10 to HH20 fate map. **(A)** HH10 fate map and HH20 regions, 4D model. **(B)** HH20 region labels. Scale bars - 250 μm.

**Fig. S8.**
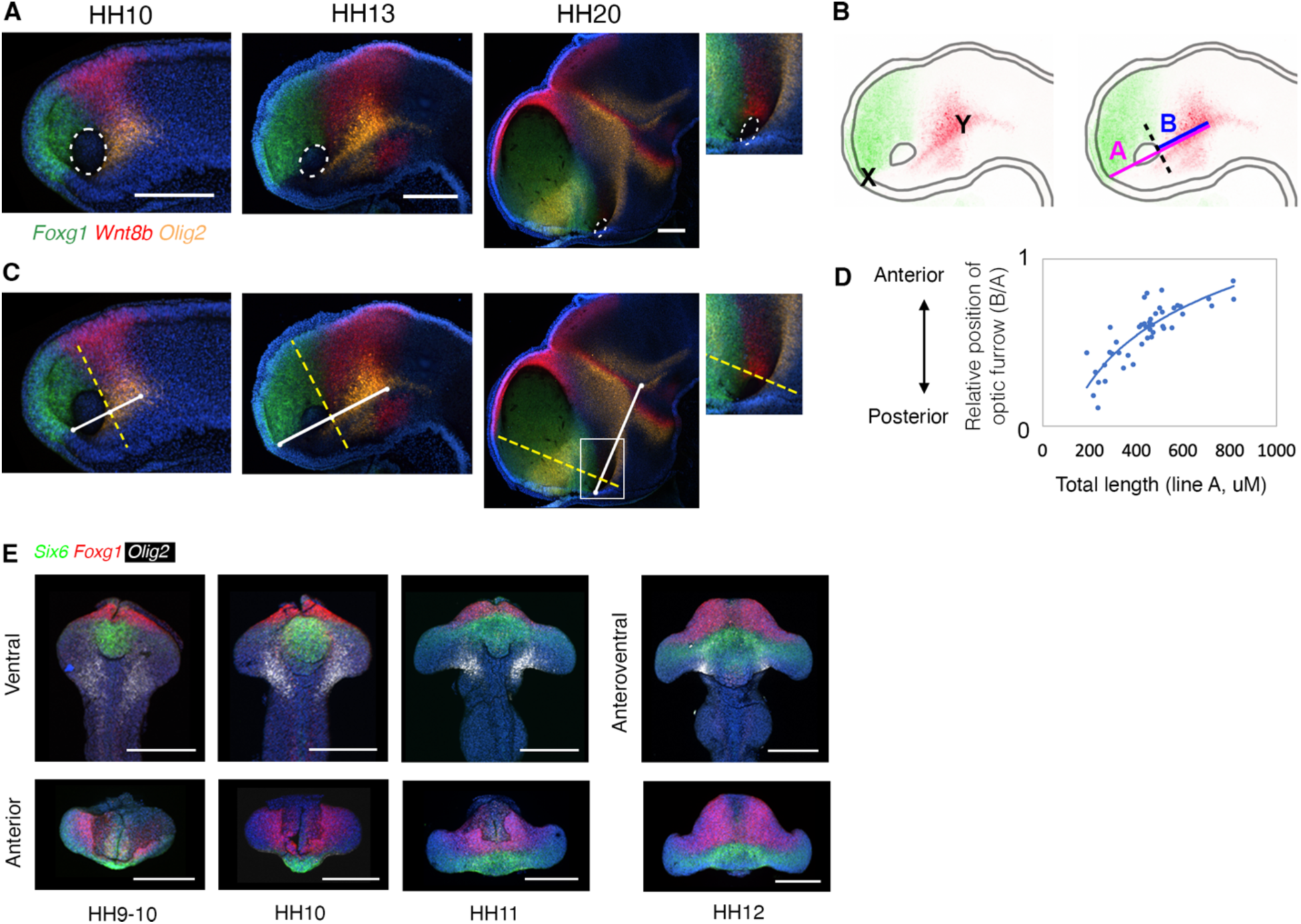
Changing dimensions of the embryonic forebrain. **(A-D)** The optic stalk draws down to a small anterior-ventral opening. (A, C) Lateral, hemisected view of HH10-20 chick brains following HCR for regional markers. Dotted line in (A) approximates the position of the retinal field at HH10 and optic stalk opening at HH13-HH20. (B-C) Positions of markers/lines used to measure relative A-P position of the posterior optic vesicle/optic stalk, as presented in (D) - see methods for details. **(E)** Dissected neuroepithelia of HH9-12 chicks processed for expression of *SIX6*, *FOXG1* and *OLIG2* by HCR. Scale bars - 250 μm.

**Fig. S9.**
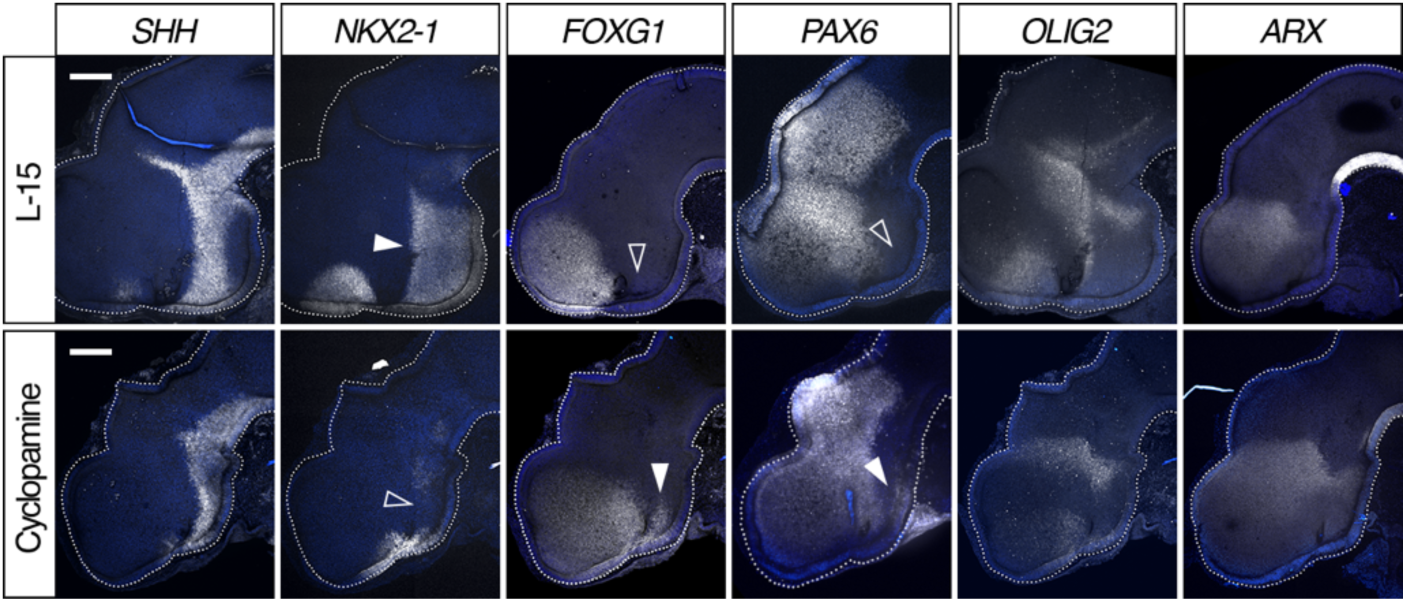
Dorsalisation of the hypothalamus. HH17-18 hemisected chick heads (internal view) after treatment with L-15 control medium (top) or cyclopamine (bottom) at HH10. HCR *in situ* for regional markers displayed in white. After cyclopamine treatment, the tuberal hypothalamus is reduced in size, *SHH* expression is reduced and *NKX2.1* hypothalamic expression is almost absent. *FOXG1* remains in the telencephalon and is also expanded into the anterior hypothalamus. *PAX6* is also ectopically expressed in the ventral tuberal hypothalamus. *OLIG2* or *ARX* persist in the posterior hypothalamus. Closed arrowheads point to regions of expression; open arrowheads to regions where expression cannot be detected. Scale bars - 250 μm.

**Fig. S10.**
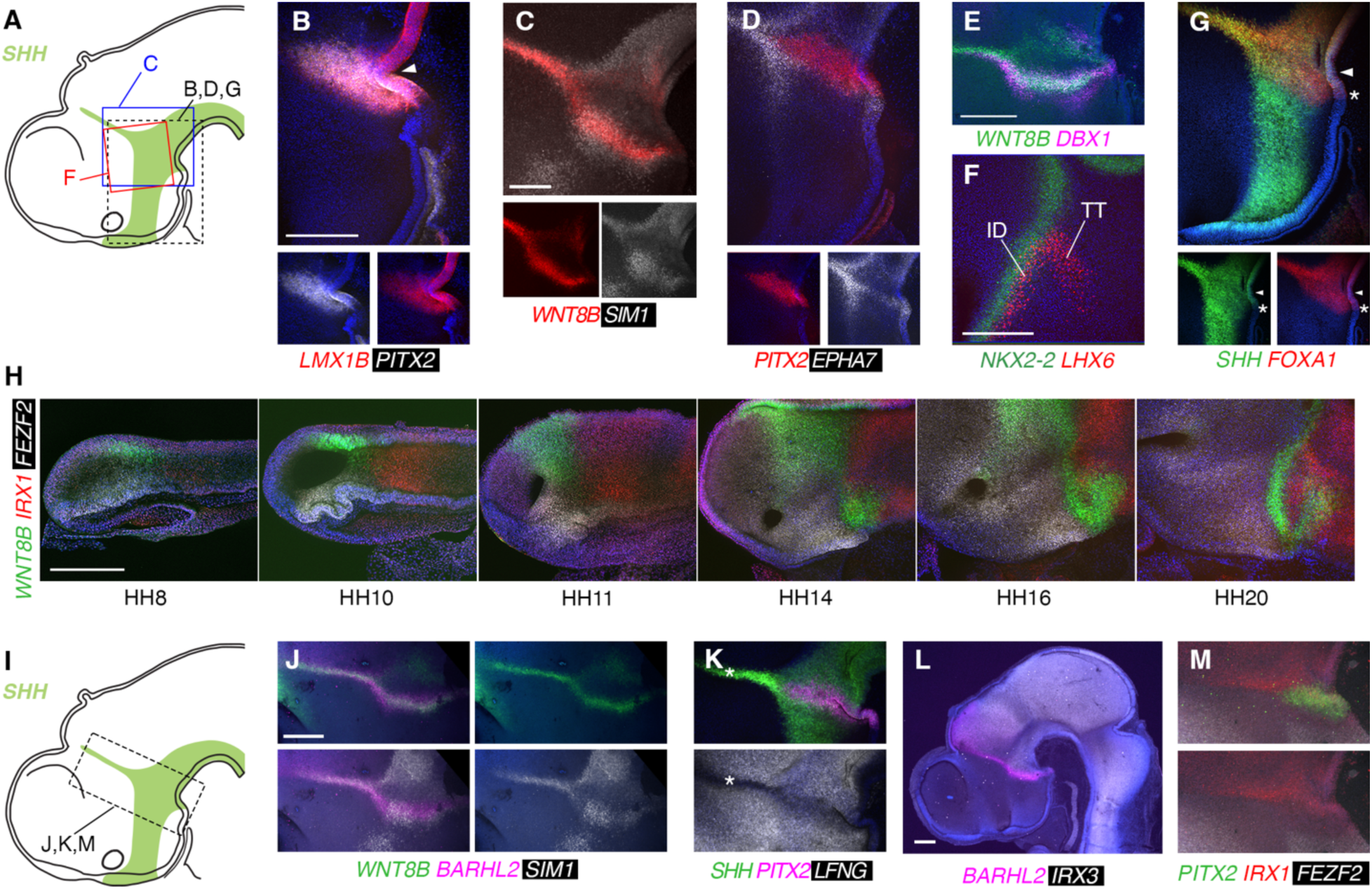
HCR analysis of chick posterior hypothalamus. **(A)** Regions shown in B-G. **(B-G)** HCR analysis of gene expression patterns in HH20 hemisected chick heads. Arrowhead in (B) and (G) indicates flexure at end of *ARX^+ve^* FP. Asterisk in (G) indicates end of *SHH*/*FOXA1^+ve^* FP. **(H)** Developmental series of HH8-HH20 chicken forebrains analysed by HCR. Hemisected lateral views with the hypothalamus oriented similarly; HH20 example is therefore rotated approximately 90 degrees clockwise compared to (B-G). **(I)** Boxed area indicates region shown in (J-K, M). **(J-M)** HCR analysis of gene expression patterns in HH20 hemisected chick heads. Scale bars - 250 μm.

**Fig. S11.**
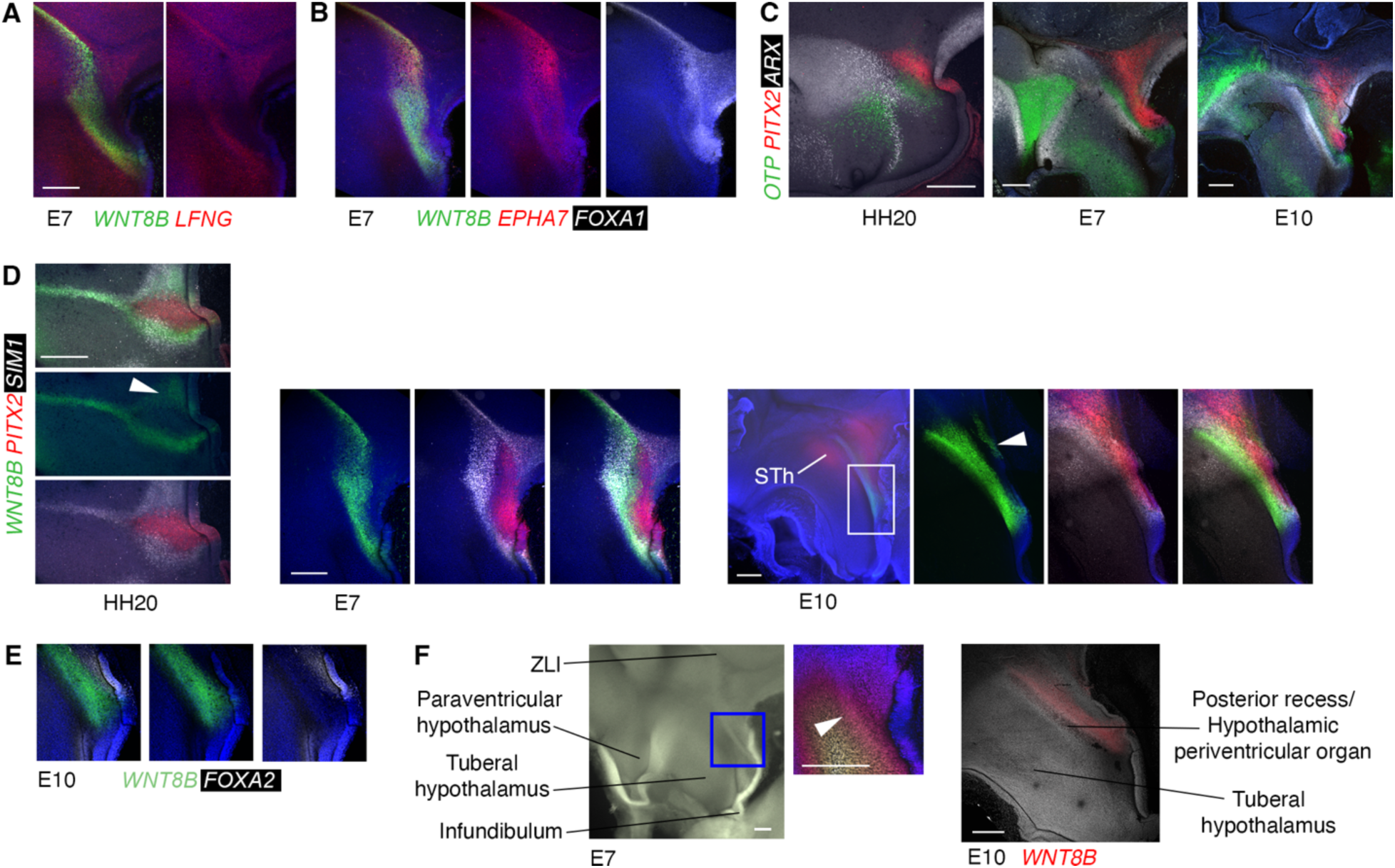
HCR analysis of chick posterior hypothalamus, HH20 - E10. **(A, B)** E7 chicken ZLI and posterior hypothalamus processed by HCR. **(C, D)** Morphological comparisons of HH20, E7 and E10 chicken hypothalamus, HCR *in situs*. Boxed area in (D) (E10) expanded to right. Arrowheads - *WNT8B* expression posterior to *PITX2*^+ve^ domain. **(E**) HCR on E10 sample showing FP *FOXA2* expression. **(F)** Morphological ridge/fold in the E7 and E10 hypothalamus. Boxed area in E7 shown expanded and coloured by z-position to highlight morphology. Same embryo as in (A), arrowhead - ridge. Scale bars - 250 μm.

**Fig. S12.**
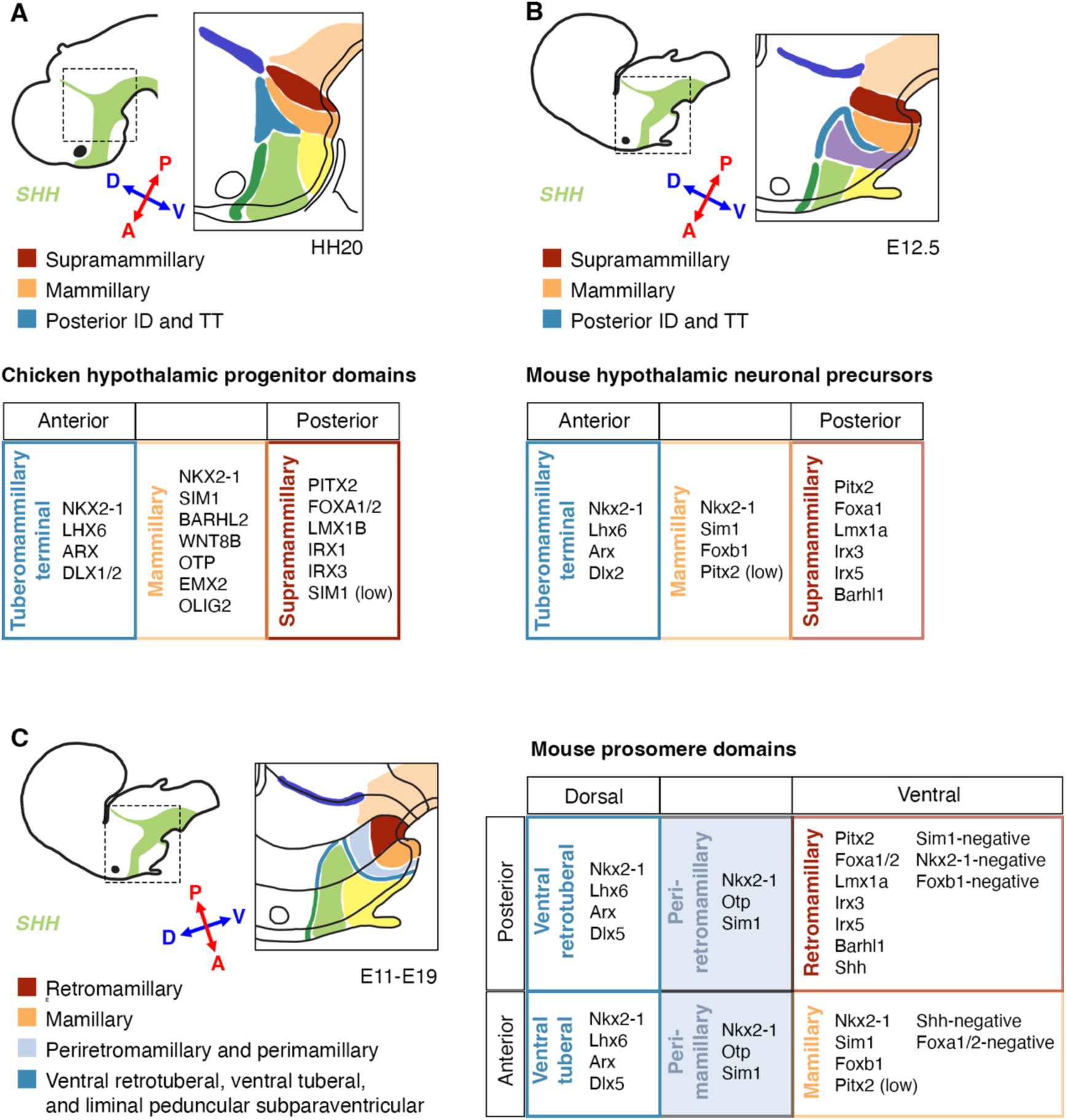
Descriptions of hypothalamic progenitor regions in chicken and mouse. **(A-C)** Hypothalamic progenitor domains, axes, and regional marker genes as described in different species and by different authors. **(A)** HH20 chicken hypothalamic progenitor domains as described in this study and (*7*), **(B)** E12.5 mouse hypothalamic neuronal precursors, as described in (*8*, *10*) **(C)** Prosomere domains as described for E11-E19 mouse (*4*, *6*, *23*, *29*). Major progenitor regions are colour coded for ease of comparison, and selected distinguishing marker genes are provided in the tables. Note that the prosomere mamillary, perimamillary and periretromamillary domains together approximately correspond to the mammillary domains in (A) and (B).

---

**Movie S1.**

**Fate Map, chicken neural tube, Hamburger-Hamilton (HH) stages 10 to 20**

**Movie S2.**

**‘Digital dye’ spots on our model recapitulate key growth patterns seen *in vivo***

**Movie S3.**

**Growth patterns shaping the anterior hypothalamus and the dorsal forebrain anterior to the ZLI**

**Movie S4.**

**Growth patterns shaping the posterior hypothalamus and the dorsal forebrain posterior to the ZLI**

